# Chemical Proteomics Reveals Protein Tyrosination Extends Beyond the Alpha-Tubulins in Human Cells

**DOI:** 10.1101/2022.07.02.498566

**Authors:** Dmytro Makarov, Pavel Kielkowski

**Affiliations:** LMU München, Department of Chemistry, Institute for Chemical Epigenetics – Munich (ICEM), Würmtalstrasse 201, 81375, Munich, Germany

**Keywords:** Tyrosination, Microtubules, Chemical proteomics, Neuronal differentiation

## Abstract

Tubulin detyrosination-tyrosination cycle regulates the stability of microtubules. Thus far described on α-tubulins, the tyrosination level is maintained by a single tubulin-tyrosine ligase (TTL). However, the precise dynamics and tubulin isoforms which undergo (de)tyrosination in neurons are unknown. Here, we exploit the substrate promiscuity of the TTL to introduce an *O*-propargyl-L-tyrosine in neuroblastoma cells and neurons. Mass spectrometry-based chemical proteomics in neuroblastoma cells using the *O*-propargyl-L-tyrosine probe revealed previously discussed tyrosination of TUBA4A, MAPRE1, and other non-tubulin proteins. This finding was further corroborated in differentiating neurons. Together we present the method for tubulin tyrosination profiling in living cells. Our results show that detyrosination-tyrosination is not restricted to α-tubulins with coded C-terminal tyrosine and is thus involved in fine-tuning of the tubulin and non-tubulin proteins during neuronal differentiation.

---

Microtubules (MTs) are composed of α- and β-tubulin heterodimers and are essential for function and stability of the cellular cytoskeleton. The defined MTs composition is critical for intracellular transport, mechanical resistance, mitosis and migration.^[1]^ Both α- and β-tubulin are encoded in the human genome in multiple isotypes, which have been observed to be tissue and cell type specific.^[2]^ The heterogeneity of MTs is further extended by numerous post-translational modifications (PTMs) including acetylation, (poly)glutamylation, (poly)glycylation, (poly)amination and tyrosination together called the tubulin code (Figure 1A).^[3]^ The majority of these PTMs are concentrated on the disordered tubulin C-terminus. In most of the α-tubulins, the encoded C-terminal amino acid is tyrosine, which can be cleaved by VASH-SVBP complex or most recently discovered MATCAP carboxypeptidase (Figure 1B).^[4]^ The terminal tyrosine is restored in translation independent manner by tubulin-tyrosine ligase. Regulation of the tyrosination statutes fine-tunes the stability of α- and β-tubulin heterodimer and MTs.^[5]^ The stable MTs induced by paclitaxel show an increased amount of detyrosinated α-tubulin, while tubulin heterodimers and unstable MTs are characterized by tyrosinated α-tubulin.^[5]^ However, the tyrosinated tubulins are not required for MTs polymerization nor MTs detyrosination is sufficient for their stabilization. The physiological relevance was demonstrated on TTL knockout mice, which die perinatally and display dysregulated development of neuronal networks.^[6]^ The upregulated MTs detyrosination was linked to failing hearts in patients with ischaemic cardiomyopathy.^[7]^ On the cellular level, MTs detyrosination is critical for directional transport of chromosomes and governance of interaction between MTs and microtubule-associated proteins (MAPs).^[7]^ Overall tubulin tyrosination was previously analyzed by cell pretreatment with cycloheximide (CHX) to block the protein synthesis and subsequent addition of tritiated tyrosine analogue to track the tyrosine incorporation by radioactivity.^[8]^ Nowadays, the tubulin (de)tyrosination status is usually identified by PTM-specific antibodies.^[8]^ However, these do not distinguish individual tubulin isotypes. On the other hand, liquid chromatography (LC) with mass spectrometry (MS)-based detection of tyrosinated and detyrosinated peptide fragments is not feasible due to the very high complexity of C-terminal peptide PTMs.

**Figure 1.**
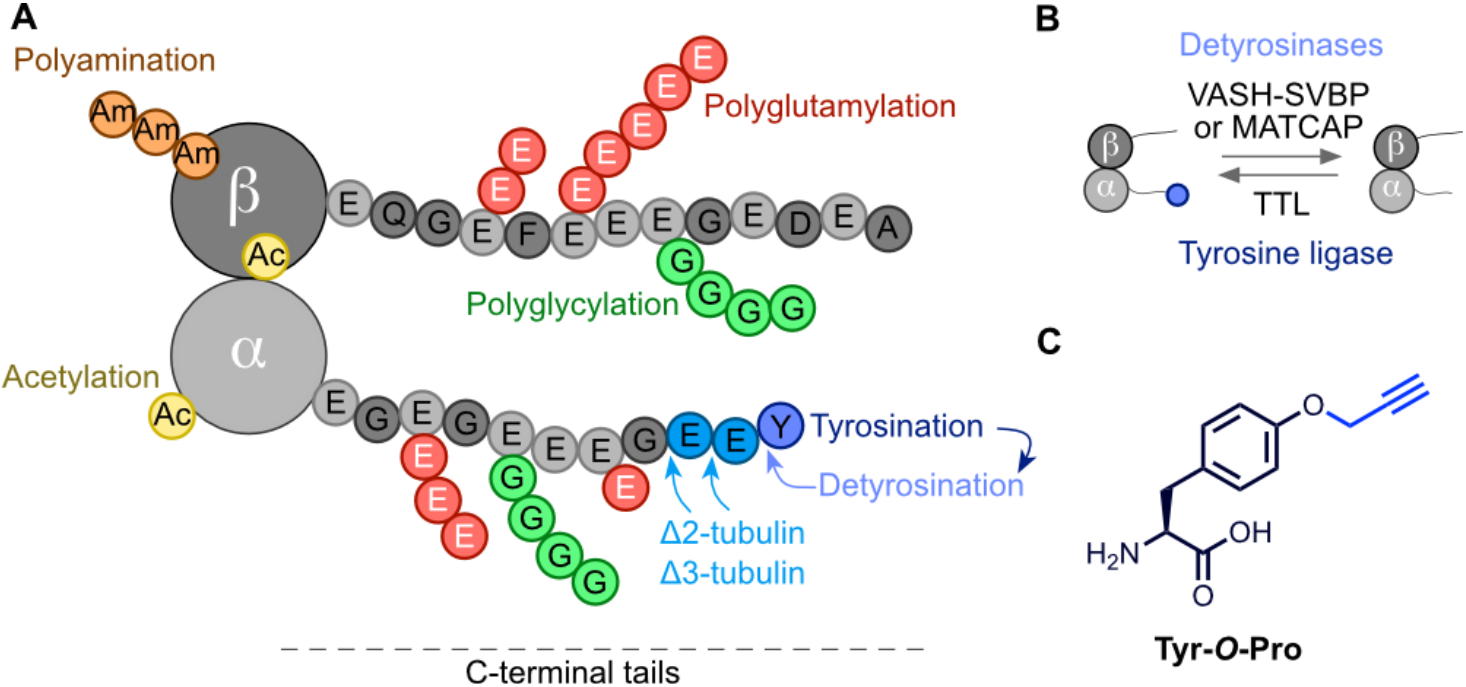
Microtubules detyrosination and tyrosination cycle. A) Tubulin code overview. B) Enzymes involved in detyrosination and tyrosination. C) Structure of the **Tyr-O-Pro** probe.

Here, we report an MS-based chemical proteomics approach to decipher the composition of post-translationally tyrosinated proteins in living cells. The approach is based on the low substrate selectivity of the TTL. The TTL’s active site was shown to be able to accommodate various tyrosine analogues.^[9]^ In contrast, modified tyrosines are poor substrates for the translation machinery. Thus, avoiding unspecific labelling of bulk proteins.

To initiate the study, we have designed and synthesized the *O*-propargyl-L-tyrosine (**Tyr-*O*-Pro**, Figure 1C). This was synthesized in three steps from L-tyrosine (Figure S1). First, the cytotoxicity of the **Tyr-*O*-Pro** was evaluated on SH-SY5Y neuroblastoma cells to show no toxicity up to 2 mM final concentration of the probe in cell culture media (Figure S2). Second, a series of in-gel experiments were carried out to optimize the labelling efficiency and to test the probe’s fidelity (Figure 2A). The **Tyr-*O*-Pro** treatment times of SH-SY5Y cells were optimized. To evaluate the extent of **Tyr-*O*-Pro** incorporation, the probe-treated cells were lysed and reacted under copper-catalyzed azide-alkyne cycloaddition (CuAAC) conditions with rhodamine-PEG_3_-azide (TAMRA-N_3_). The labelled proteins were separated using sodium dodecyl sulfate-polyacrylamide gel (SDS-PAGE) electrophoresis and rhodamine fluorescence was scanned. The successful protein labelling was observed already between 8 to 16 hours after addition of the 300 μM **Tyr-*O*-Pro** into cell culture media but further increased for a total of two days (Figure 2B). Interestingly, the strongest fluorescence band exhibiting time-dependent labelling was observed at around 50 kDa, suggesting the labelling of tubulins. In parallel, probe concentration was tested to reveal that already 200 μM final concertation provides bright fluorescent bands (Figure S3). Next, to confirm the translation-independent incorporation of **Tyr-*O*-Pro** in proteins, the SH-SY5Y cells were pre-treated with CHX to block the protein synthesis before **Tyr-*O*-Pro** probe addition (Figure 2C). Indeed, we observed only a slight decrease in fluorescence intensity likely due to an overall decrease in protein/tubulin amount. This confirms the introduction of **Tyr-*O*-Pro** as protein PTM rather than via ribosomal protein synthesis.

**Figure 2.**
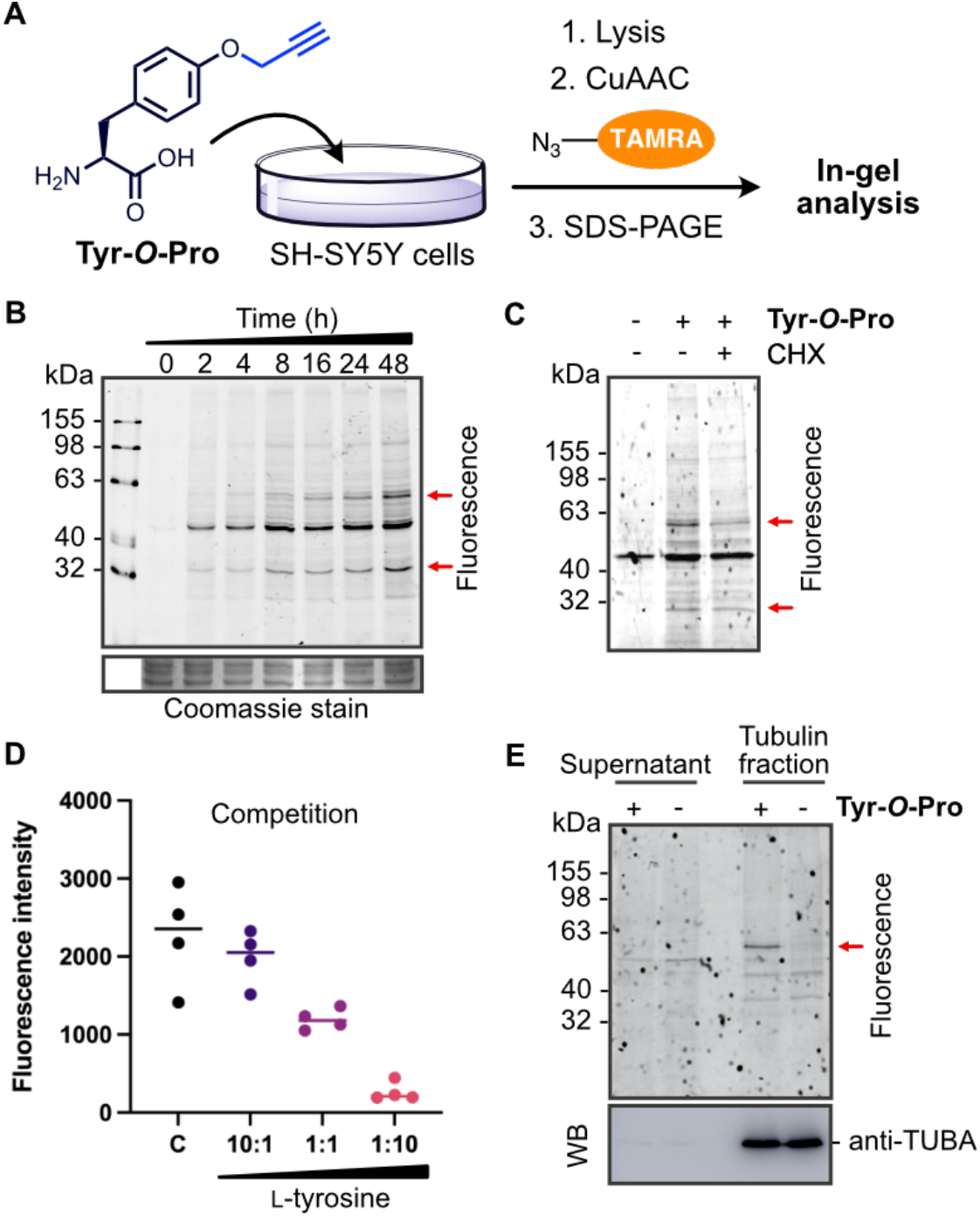
In-gel analysis of **Tyr-O-Pro** protein metabolic labelling in SH-SY5Y cells. A) The overall metabolic labelling strategy. B) Time-dependent labelling using the **Tyr-O-Pro** probe (300 μM). Cells were harvested at indicated time points after addition of the probe. C) The cells were pre-treated with CHX for 30 min before addition of the **Tyr-O-Pro** probe and incubation for 24 h. This excludes the incorporation of the probe via ribosomal translation machinery. D) The cells were treated with a constant final concentration of the **Tyr-O-Pro** probe and further supplemented with the increasing amount of L-tyrosine as indicated. E) In-gel fluorescence of the isolated tubulin fraction reveals a single specific band at around 55 kDa. Abbreviation: C: control, only **Tyr-O-Pro** probe treated cells. WB: Western blot.

Furthermore, a competition experiment between the probe and natural tyrosine was performed to test the probe’s fidelity. Indeed, the probe labelling was clearly diminished with an increasing amount of natural tyrosine (Figure 2D and Figure S4). Next, the tubulin fraction was isolated by taxol-induced depolymerization-polymerization of MTs from **Tyr-*O*-Pro** treated SH-SY5Y cells.^[8b,10]^ The single probe-specific fluorescent band was observed in the tubulin fraction corroborating the fidelity of the probe (Figure 2E). Tubulin fraction isolation was tested by western blot using the anti-TUBA antibody. Finally, the turnover of the **Tyr-*O*-Pro** probe was examined by treatment of the SH-SY5Y cells with the probe followed by cell culture media exchange without the probe. Surprisingly, we observed a rather slow replacement of incorporated propargyl tyrosine by natural tyrosine, only after 24 h, there was no observable band present (Figure S5). Together, these experiments support the efficient incorporation of **Tyr-*O*-Pro** in tubulins as PTM.

To decipher the composition of tubulin isoforms labelled by the **Tyr-*O*-Pro** probe we continued with MS-based chemical proteomics. Recently, we have established an efficient chemical proteomics enrichment approach called SP2E.^[11]^ The SP2E workflow uses the carboxylate-modified magnetic beads to clean up the proteins after the click chemistry with biotin-N_3_. In the next step, the proteins are eluted from the carboxylate magnetic beads and transferred on streptavidin-coated magnetic beads. The biotin-labelled proteins are enriched and digested by trypsin. The resulting peptide mixtures corresponding to the enriched probe-modified proteins are collected and analyzed via LC-MS/MS. In parallel, the same procedure is carried out with control cells to be able to abstract the background, which is composed of non-specifically enriched proteins (Figure 3A). We have utilized the SP2E workflow to enrich **Tyr-*O*-Pro** labelled proteins from SH-SY5Y cells in quadruplicates using the data-independent acquisition (DIA).^[12]^ Indeed, analysis of resulting MS spectra by DIA-NN^[12]^ via label-free quantification (LFQ)^[12]^ confirmed the significant enrichment of α-tubulin isoforms including TUBA1C and TUBA4A and TUBGCP5 (Figure 3B). To gain more confidence in the identification of the highly similar tubulin isoforms, we have revised the peptides assigned to the tubulin isoforms together with corresponding MS2 spectra (Table S1). This analysis showed that at least 2 unique peptides were found for each α-tubulin isoform. The TUBA4A isoform does not encode the C-terminal tyrosine resembling the detyrosinated tubulin when translated. However, TUBA4A was speculated to be possibly tyrosinated, which is now corroborate by our results. Surprisingly, several non-tubulin proteins were found significantly enriched as well. This group contains the microtubule-associated protein RP/EB family member 1 (MAPRE1 also called EB1) a marker of the microtubule plus-end, which regulates the dynamics of the microtubule cytoskeleton.^[13]^ MAPRE1 is involved in a mitotic spindle positioning and recruiting the CLIP170 to MTs (+)-end.^[13]^ Importantly, sequence analysis of this 30 kDa protein contains the α-tubulin-like C-terminus coding the terminal tyrosine adjacent to two glutamates (PQEEQ**EEY**). This would suggest a potential detyrosination-tyrosination cycle as discussed in the literature.^[14]^ The retrospective analysis of the in-gel fluorescence labelling shows clear time-dependent labelling of a protein at around 30 kDa (Figure 2B). Some other non-tubulin proteins were significantly and consistently enriched (as well in the following experiments in iNGNs) such as MARCKS, DCTN3 and LSM6. From these, the MARCKS and DCTN3 are proteins with a role in cytoskeleton organization. In parallel, the **Tyr-*O*-Pro** probe treated lysates were processed by SP3^[15]^ for whole proteome MS analysis to exclude the translation-dependent stochastic incorporation of the probe into proteins. The resulting MS spectra were searched for the presence of *O*-propargyl modified tyrosine in identified peptides. In both control and **Tyr-*O*-Pro** treated cells on average only 42.5 and 33 modified proteins were found from average of 38606 and 37291 total peptides respectively (Table S2). Thus, suggesting the **Tyr-*O*-Pro** is indeed incorporated post-translationally. With the established **Tyr-*O*-Pro** labelling and MS-based chemical proteomics workflow, we moved towards the application of the method to determine the dynamics of protein tyrosination during neuronal differentiation.

**Figure 3.**
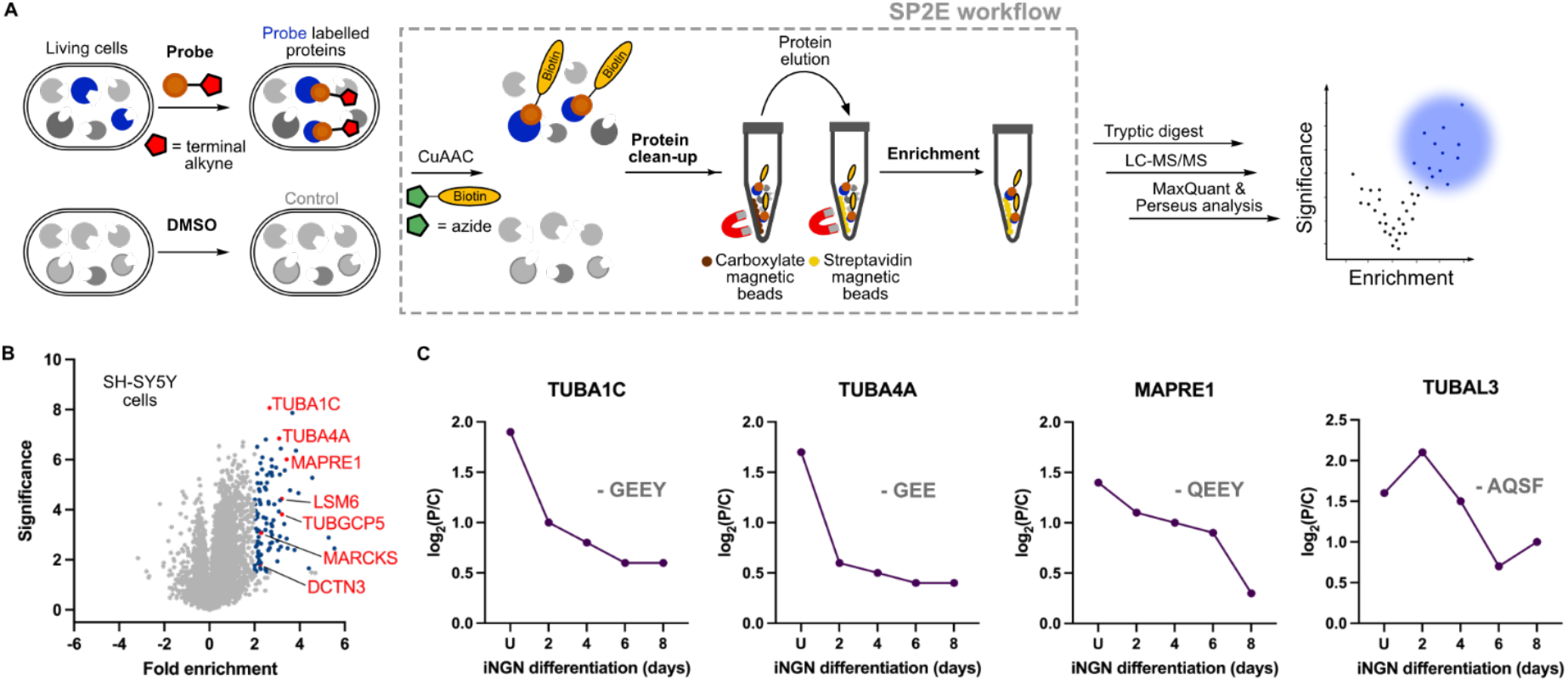
Mass spectrometry-based chemical proteomics of SH-SY5Y and iNGN cells uncovered the scope of protein tyrosination. A) Overview of MS-based chemical proteomics workflow using the SP2E. B) Volcano plot visualizing the enrichment of **Tyr-*O*-Pro** labelled proteins from SH-SY5Y cells; Blue dots are significantly enriched proteins with at least 4-fold enrichment. *n* = 4, fold enrichment (log_2_(**Tyr-*O*-Pro**/control), significance (-log_10_(*p*-value)). C) Fold change of tyrosination on selected proteins during iNGNs neuronal differentiation showing tyrosination/detyrosination cycle of TUBA4A and MAPRE1. The isoform encoded C-terminal sequences are added for comparison

The neuronal cells are characterized by their strong polarization of the cell body containing dendrites, axons and synapses. The neuronal cytoskeleton is responsible for the maintenance of this polarization, it supports the neuronal migration during cortex development and provides avenues for cellular trafficking.^[16]^ The composition of tubulin isoforms is known to play a role during neuronal differentiation. However, it was previously not possible to link the tubulin isoforms with corresponding tyrosination status. We applied the **Tyr-*O*-Pro** probe to determine protein tyrosination status during neuronal differentiation of human-induced pluripotent stem cells (hiPSCs). To streamline this process, we leveraged from fast (4 days) differentiation of hiPSCs engineered with doxycycline-inducible Neurogenin-1 and -2 cassette (iNGNs).^[17]^ To test the feasibility of our approach in iNGNs, they were treated with **Tyr-*O*-Pro** at four different time points during differentiation into mature neurons (Figure S6). Similar to SH-SY5Y cells, the fluorescent band at around 50 kDa likely corresponding to the tubulins was present (Figure S6). In addition, we observed a clear change in fluorescence intensity of the 30 kDa band during iNGNs differentiation (Figure S6).

Thus, we have proceeded with the SP2E enrichment of **Tyr-*O*-Pro** labelled proteins using the trifunctional linker (containing 5/6-TAMRA-N_3_-biotin moieties) instead of the biotin-N_3_. In contrast to the previous MS experiments, enriched proteins were eluted from streptavidin magnetic beads using the SDS-PAGE loading buffer. Fluorescence imaging of the gel showed the labelling in a region around 55 and 30 kDa. The subsequent western blot of these enriched proteins and staining with the anti-MAPRE1 antibody showed the presence of MAPRE1 protein pool in probe-treated cells 2 days after doxycycline-induced neuronal differentiation (Figure S7). Next, the MS-based chemical proteomics was performed using the small-scale 96-well plate format SP2E workflow starting with 100 μg protein. In total, we have collected the cells at five time points during the iNGNs differentiation and maturation. We have observed a decrease in overall tubulin tyrosination on TUBA1C, TUBA4A and MAPRE1 during the neuronal differentiation (Figure 3C and Figure S8). Moreover, tubulin alpha chain-like 3 (TUBAL3) protein was identified showing a different pattern. Despite the high sequence similarity with α-tubulins (>75%), the TUBAL3 C-terminus lacks the -EEY motive. Several other non-tubulin but cytoskeleton-associated proteins were significantly enriched suggesting their tyrosination including dynactin subunit 3 (DCTN3), alpha-tubulin N-acetyltransferase 1 (ATAT), MARCKS-related proteins MARCKS and MARCKSL1. Surprisingly, a cytosolic U6 snRNA-associated Sm-like protein LSm6 (LSM6) was consistently enriched at all time points during the iNGNs differentiation (Figure S8). SP2E enrichment experiment in iNGNs was complemented by whole proteome analysis. This confirmed the identity of the neuronal cells and also aided to estimate trends in tyrosination stoichiometry and thus dynamics. In general, there is a strong increase in total tubulin isoforms expression (Figure S9). The same trend was found for the MAPRE1 explaining the above-described observation using the western blot as the read-out. It was possible to detect MAPRE1 after enrichment with anti-MAPRE1 antibody only in two-day differentiated iNGNs, because of the sufficient absolute amount of tyrosinated MAPRE1 in the lysate. While the MAPRE1 tyrosination (fold enrichment) mildly decreases after two days, there is at the same time a dramatic increase in MAPRE1 protein expression (Figure 3C and Figure S9).

The tyrosination status of the proteins correlates well with the expression of the TTL, which is lower than that of carboxypeptidases responsible for the tyrosine removal (VASH1 and MATCAP) (Figure S10). Together, these experiments provide evidence of significant changes in tyrosination status during neuronal differentiation on tubulin and non-tubulin proteins.

To visualize the probe distribution and incorporation within the SH-SY5Y cells, the **Tyr-*O*-Pro** probe was used for fluorescence imaging (Figure 4). Control or probe-treated cells were washed to remove excess of the probe and incubated with TAMRA-N_3_ under CuAAC conditions. The strong labelling in the probe-treated cells was observed, with a negligible background in control. Colocalization with TUBA and MAP2 showed partial overlap, which is in the line with most tyrosinated tubulins being present in cells as dimers, which are not polymerized in microtubules.

**Figure 4.**
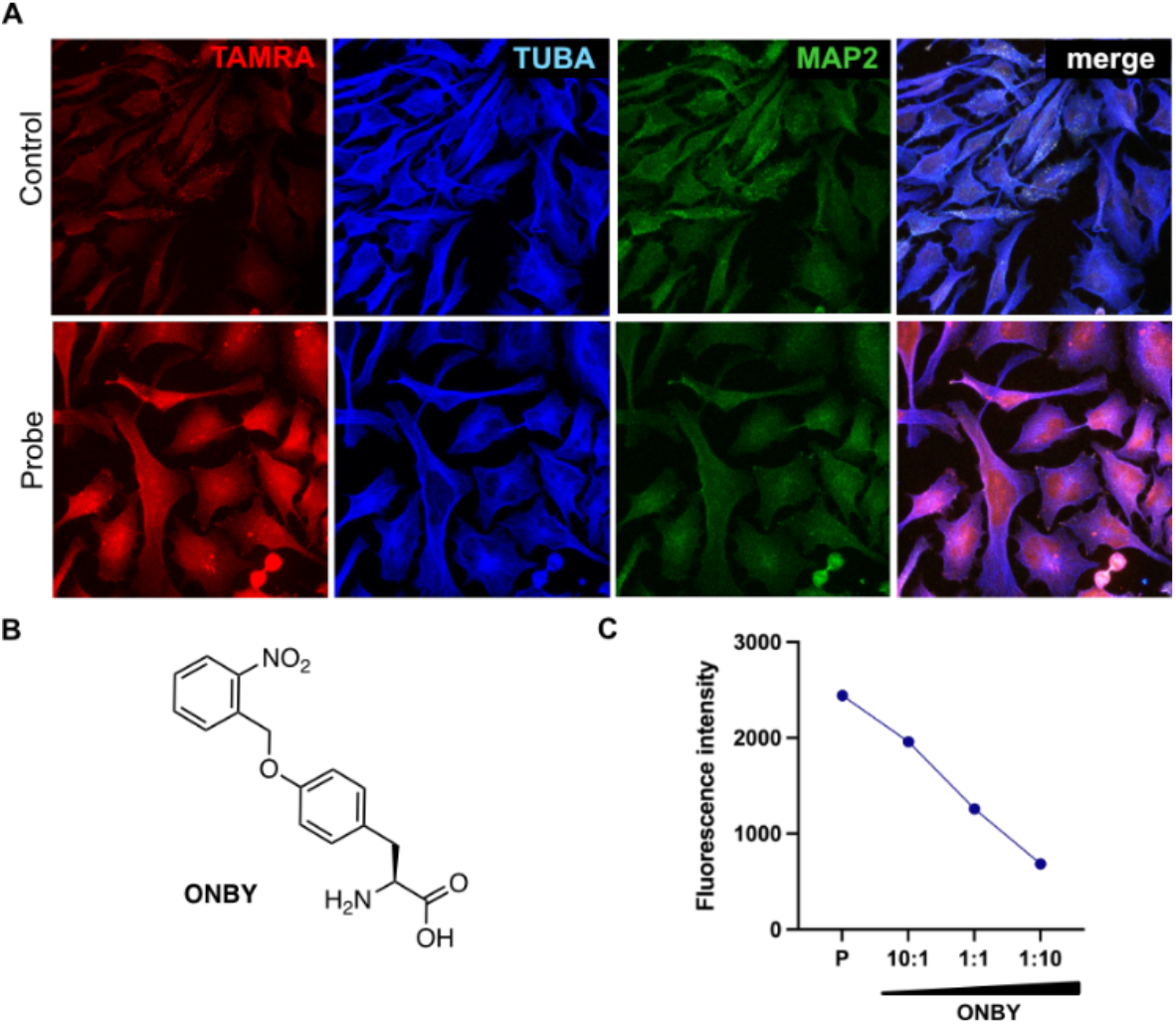
Fluorescence imaging and incorporation of sterically hindered tyrosine derivative. A) Fluorescence imaging of **Tyr-*O*-Pro** treated and control SH-SY5Y cells. B) ONBY structure. C) In-gel analysis of the competition experiment between **Tyr-*O*-Pro** and ONBY in SH-SY5Y cells. The fluorescence band at around 55 kDa was quantified. P – only **Tyr-*O*-Pro** treated cells.

Finally, we were interested in whether it is possible to further utilize the low substrate selectivity of TTL for introduction of other functional groups in living cells. The selected *O*-(2-nitrobenzyl)-L-tyrosine (**ONBY**) was added together with **Tyr-*O*-Pro** to the cells in different ratios. The final concentration of **Tyr-*O*-Pro** was kept constant. This setup resulted in the competition between the two unnatural tyrosines, which was then analyzed after CuAAC with TAMRA-N_3_ and fluorescence scanning of the SDS-PAGE (Figure 3D and Figure S11). The MTs incorporated ONBY tubulin might be further used to probe the MTs’ interactions with MAPs. The nitrobenzyl residue can be removed by UV-light irradiation, which would enable to modulate the tyrosination function in the living cells.^[18]^

In summary, a straightforward approach for chemical proteomic analysis of protein tyrosination in living cells based on the metabolic labeling using the **Tyr-*O*-Pro** probe was developed. Our method enables for the first time to profile tyrosinated tubulin isoforms during neuronal differentiation and strongly suggests, that tyrosination is not restricted to α-tubulins. We have confirmed disputed tyrosination of microtubule-associated protein MAPRE1 and TUBA4A as well as on other non-tubulin proteins. Microtubules are critical for mechanical resistance of the cells and intracellular trafficking, the hallmarks of cancer research and neurodegeneration respectively. Further research in this direction enabled by the reported strategy will be carried out in our laboratory to determine the pathophysiological relevance of protein tyrosination.

## Acknowledgements

This work was funded by Liebig fellowship (Fonds der chemischen Industrie) to P.K. and LMUexcellent Junior Research Fund. We are grateful for generous support from SFB 1309 – 325871075 (Deutsche Forschungsgemeinschaft).

## Declaration of Interests

The authors declare no competing interests.

## Notes

### Competing Interest Statement

The authors have declared no competing interest.

