## Supporting Information for "Chemical Proteomics Reveals Protein Tyrosination Extends Beyond the Alpha-Tubulins in Human Cells"

#### Table of Contents

**Supplementary Figures** ... 3

**Supplementary Tables** ... 9

**Organic synthesis** ... 11

Synthesis of (*tert*-butoxycarbonyl)-*L*-tyrosine (2)[1] ... 11

Synthesis of prop-2-yn-1-yl (*S*)-2-amino-3-(4-(prop-2-yn-1-yloxy)phenyl)propanoate(4)[1] ... 11

Synthesis of (*S*)-2-amino-3-(4-(prop-2-yn-1-yloxy)phenyl)propanoic acid (Tyr-*O*-Pro, 6)[3] ... 12

Synthesis of ONBY ... 12

Synthesis of *tert*-butyl *L*-tyrosinate (7)[4] ... 12

Synthesis of *tert*-butyl (*tert*-butoxycarbonyl)-*L*-tyrosinate (8) ... 13

Synthesis of *tert*-butyl (*S*)-2-((*tert*-butoxycarbonyl)amino)-3-(4-((2-nitrobenzyl)oxy)phenyl)propanoate (9) ... 13

Synthesis of (*S*)-2-amino-3-(4-((2-nitrobenzyl)oxy)phenyl)propanoic acid (10) ... 14

**Biochemistry** ... 15

Cell lines ... 15

SH-SY5Y cell line ... 15

iNGNs cell line ... 15

Cells treatment and harvesting ... 16

Lysates preparation ... 16

Protein concentration measurement ... 16

SP2E workflow large scale ... 16

In-gel analysis of the enriched proteins ... 17

Sample preparation for MS analysis ... 17

SP2E workflow small scale ... 17

In-gel analysis ... 18

Western blot ... 18

#### Supplementary Figures

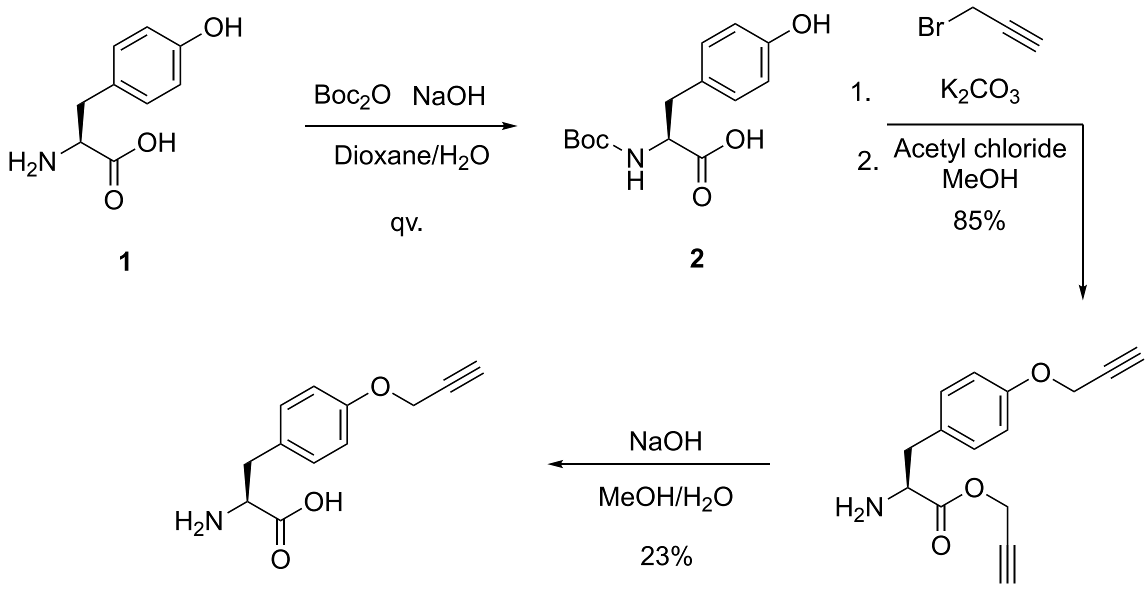

**Figure S1**. Synthesis of the **Tyr-*O*-Pro** probe. The three-step synthesis starts with L-tyrosine **1** to Boc-protected L-tyrosine **2**, which enters nucleophilic substitution reaction with subsequent detachment of Boc protection group. The ester hydrolysis of the compound **4** resulted in the desired probe **6**.

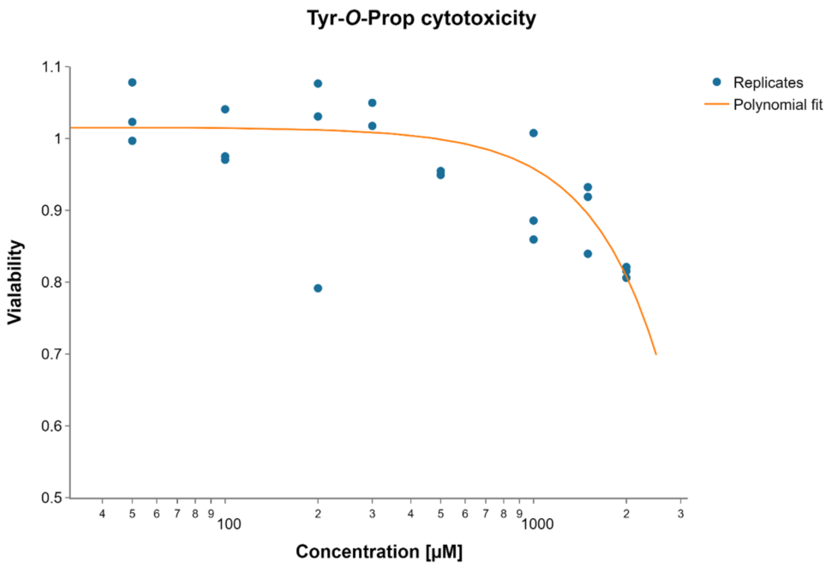

**Figure S2**. **Tyr-*O*-Pro** MTT cytotoxicity evaluation in SH-SY5Y cells. The final concentration of the probe is plotted on x-axis in log-scale; y-axis represents viability. Viability of the control group was set to 1.

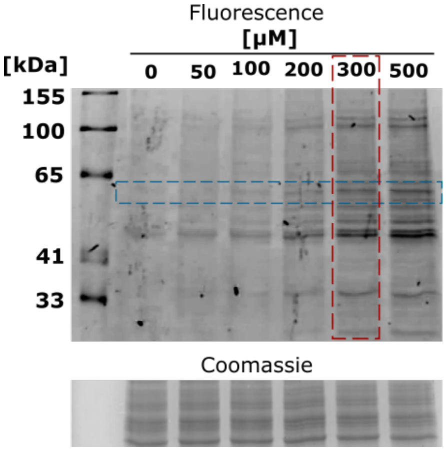

**Figure S3**. Dose-dependent in-gel fluorescence analysis of **Tyr-*O*-Pro** probe labelling in SH-SY5Y cells. Probe concentration of 0.3 mM was used within the study.

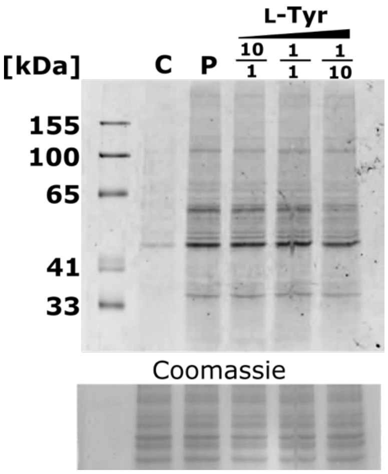

**Figure S4**. Competition experiment between natural L -tyrosine (**L-Tyr**) and the **Tyr-*O*-Pro** probe. Negative control (**C**): cells were treated with plain solvent. Positive control (**P**): cells were treated with **Tyr-*O*-Pro** probe. Other samples contain a mixture of **Tyr-*O*-Pro** and **L-Tyr** in different ratios. Concentration of **Tyr-*O*-Pro** in the samples is constant.

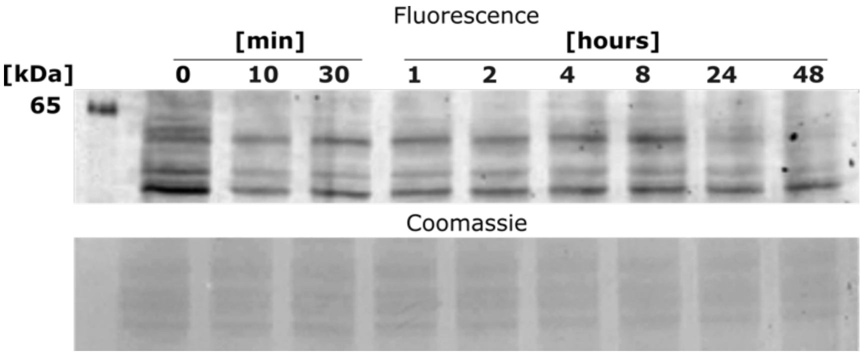

**Figure S5**. Turnover of the **Tyr-*O*-Pro** probe in SH-SY5Y cells. After one day of SH-SY5Y cells incubation with the **Tyr-*O*-Pro** probe, the medium was exchanged for the probe-free media. Cells were harvested at different time points after the media exchange.

A)

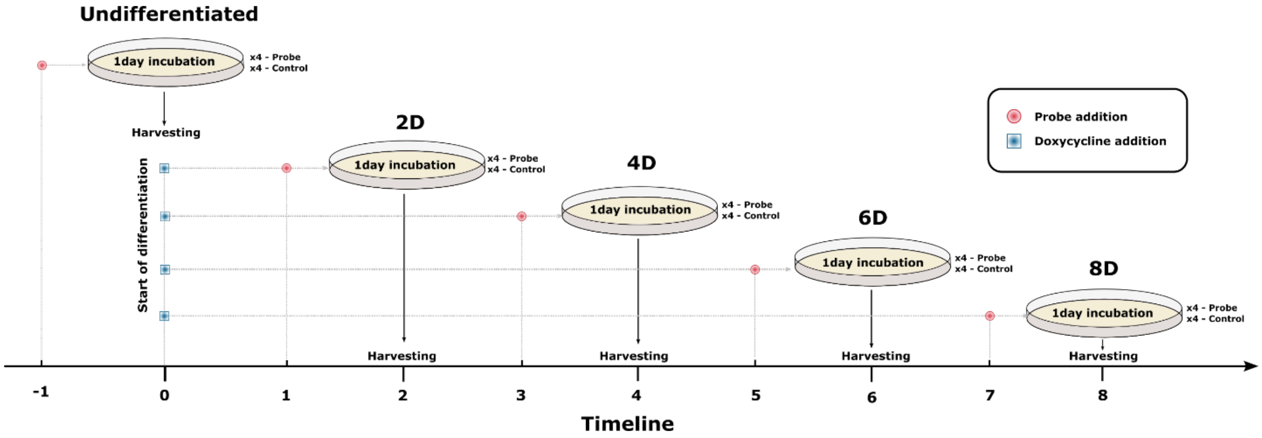

B)

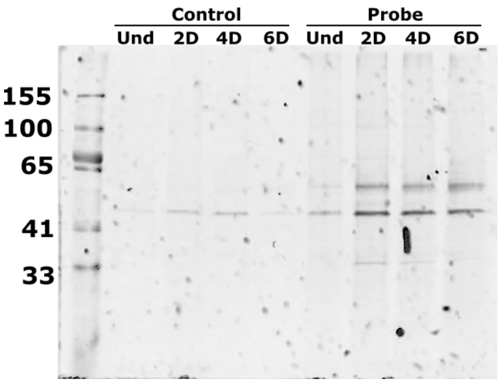

**Figure S6**. A) iNGNs probe treatment and harvesting scheme. B) In-gel fluorescence imaging from iNGNs cells upon the **Tyr-*O*-Pro** probe treatment during iNGNs differentiation. **Und**: undifferentiated iNGNs stem cells, **2D**, **4D**, **6D**, **8D**: differentiated iNGNs cells after two, four, six and 8 days after DOX addition. Control (**control**): without addition of the **Tyr-*O*-Pro** probe, and probe-treated (**probe**) samples with the **Tyr-*O*-Pro** probe in 0.3 mM concentration and 1 day incubation time.

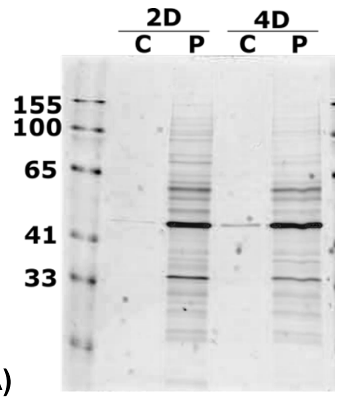

A)

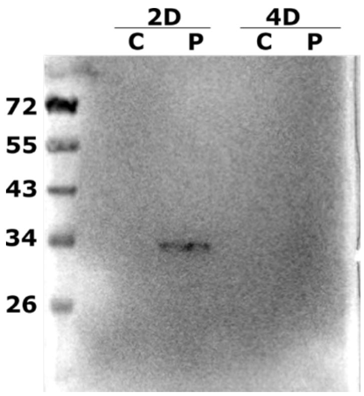

B)

**Figure S7**. The SP2E enrichment of the **Tyr-*O*-Pro** probe treated iNGNs using the trifunctional linker (TAMRA-$N_3$-biotin). **A)** In-gel fluorescence after proteins elution from the streaptavidine magnetic beads. **B)** Western blot with anti-MARPRE1 antibody. Differentiated iNGNs cells, two

(**2D**) and four (**4D**) days after differentiation started. (**C**) – control set of cells without addition of the probe; (**P**) - probe-treated cells.

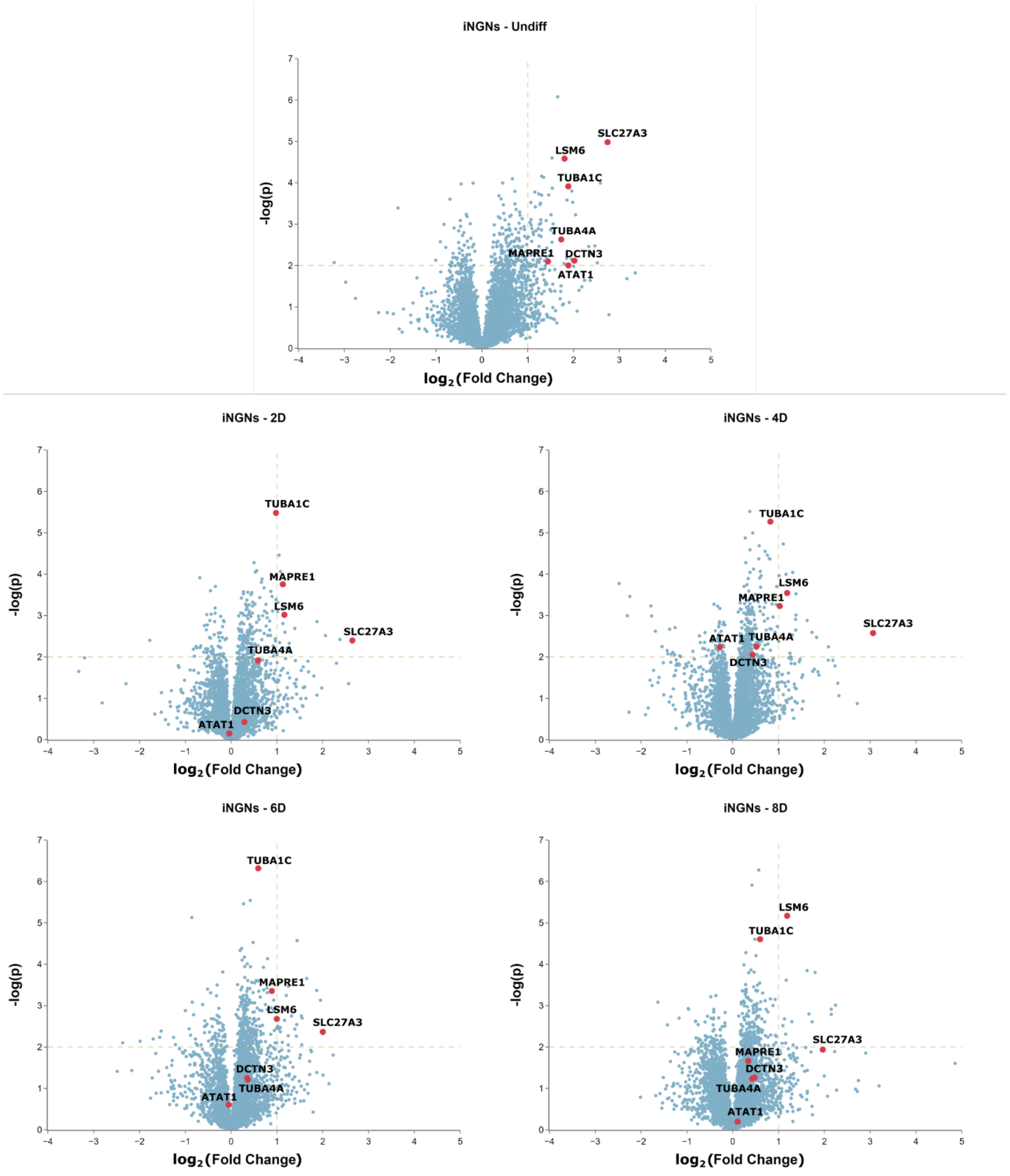

**Figure S8**. Volcano plots at different differentiation stages from iNGNs SP2E-enriched samples. The x-axis represent $log_2$-scaled relative change (**fold change**) between two conditions: probe-treated (positive values) and untreated cells (negative values). Vertical axis is a negative $log_{10}$ from p-value (statistical significance). Number of replicates for each condition = 4.

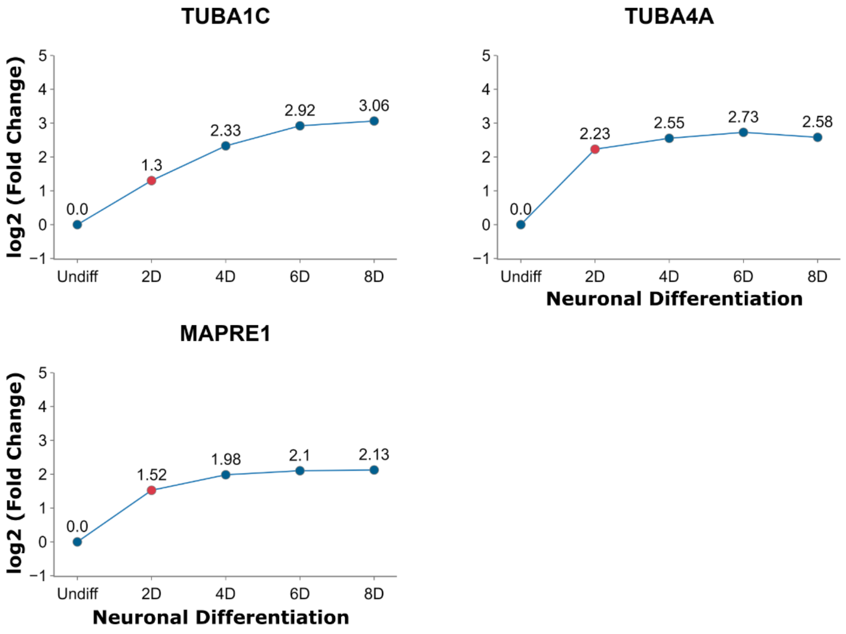

**Figure S9**. Whole proteome analysis. Graphs show protein expression changes during neuronal differentiation. Change of the protein expression is related to the first time point (undifferentiated iNGNs) and is represented in $log_2$-scale of fold change. Number of replicates for each time point = 4.

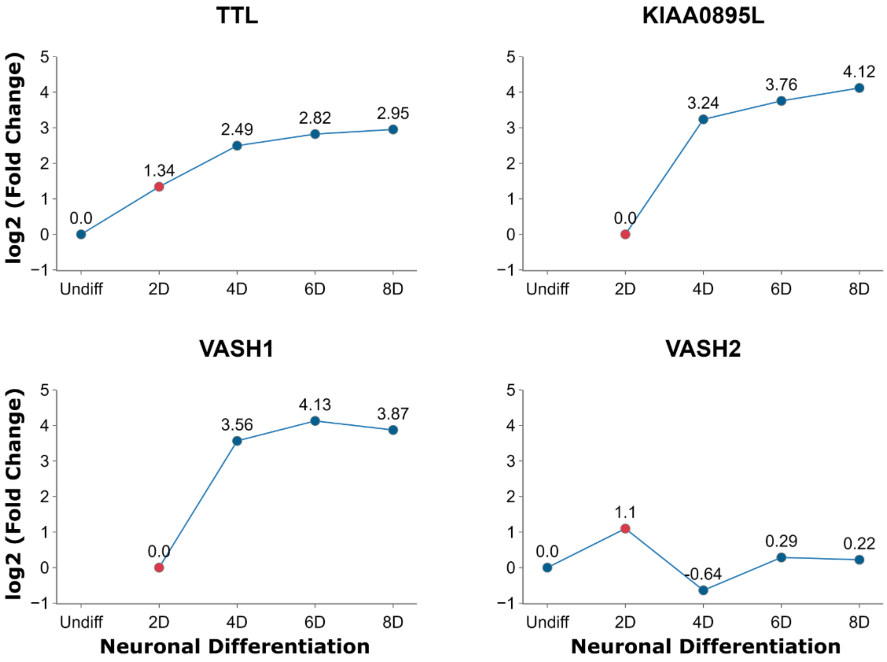

**Figure S10**. Whole proteome analysis. Graphs show protein expression change of the key detyrosination-tyrosination cycle regulators. Change of the protein expression is related to the first time point (undifferentiated iNGNs) and is represented in a $log_2$-scale of fold change. The number of replicates for each time point = 4. For VASH1 and KIAA0895L(MATCAP), no proteins were identified in undifferentiated iNGNs. 2D iNGNs serves as a first time point.

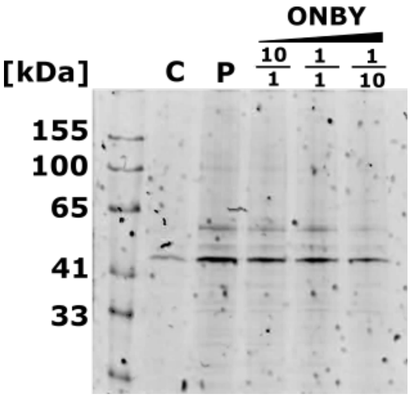

**Figure S11**. Competition experiment between *O*-(2-nitrobenzyl)-L-tyrosine (**ONBY**) and the **Tyr-*O*-Pro** probe in SH-SY5Y cells. Negative control (**C**): cells were treated with plain solvent. Positive control (**P**): cells were treated with **Tyr-*O*-Pro** probe. Other samples contain a mixture of **Tyr-*O*-Pro** and **L-Tyr** in different ratios. Concentration of **Tyr-*O*-Pro** in the competition samples is constant.

#### Supplementary Tables

**Table S1**. List of SP2E-enriched peptides found in SH-SY5Y cells samples for specific proteins.

| **TUBA1C** | | |
|---|---|---|
| Unique | Peptide sequence | Precursor charge |
| yes | AVC(Acetyl)MLSNTTAVAEAWAR | 3 |
| yes | AVC(Acetyl)M(Carbamidomethyl)LSNTTAVAEAWAR | 3 |
| | AVFVDLEPTVIDEVR | 3 |
| | DVNAAIATIK | 2 |
| | EIIDLVLDR | 2 |
| | GHYTIGKEIIDLVLDR | 3 |
| | GHYTIGKEIIDLVLDRIR | 3 |
| | GHYTIGKEIIDLVLDRIR | 4 |
| | IHFPLATYAPVISAEK | 3 |
| | LDHKFDLMYAK | 2 |
| | NLDIERPTYTNLNR | 3 |
| | QLFHPEQLITGK | 2 |
| | QLFHPEQLITGKEDAANNYAR | 4 |
| | RNLDIERPTYTNLNR | 3 |
| | TIGGGDDSFNTFFSETGAGK | 3 |
| | VGINYQPPTVVPGGDLAK | 3 |
| | VGINYQPPTVVPGGDLAKVQR | 3 |
| | YMAC(Acetyl)C(Acetyl)LLYR | 2 |
| | YM(Carbamidomethyl)AC(Acetyl)C(Acetyl)LLYR | 2 |
| **TUBA4A** | | |
| Unique | Peptide sequence | Precursor charge |
| | AVC(Acetyl)MLSNTTAIAEAWAR | 3 |
| | AVC(Acetyl)M(Carbamidomethyl)LSNTTAIAEAWAR | 3 |
| yes | AVFVDLEPTVIDEIR | 3 |
| | AYHEQLSVAEITNAC(Acetyl)FEPANQMVK | 4 |
| | AYHEQLSVAEITNAC(Acetyl)FEPANQM(Carbamidomethyl)VK | 4 |
| yes | EIIDPVLDR | 2 |
| **MAPRE1** | | |
| Unique | Peptide sequence | Precursor charge |
| | FFDANYDGK | 2 |
| | KFFDANYDGK | 2 |
| yes | KPLTSSSAAPQRPISTQR | 3 |
| yes | LEHEYIQNFK | 2 |
| yes | QGQETAVAPSLVAPALNKPK | 3 |
| yes | QGQETAVAPSLVAPALNKPK | 4 |

**Table S2**. Number of peptides found with variable modification of propargyl group (38.0156 m/z shift) on a tyrosine amino acid. SH-SY5Y cells; whole proteome samples.

| | Contr. 1 | Contr. 2 | Contr. 3 | Contr. 4 | Probe 1 | Probe 2 | Probe 3 | Probe 4 |
|---|---|---|---|---|---|---|---|---|
| Peptides with modification | 44 | 39 | 38 | 49 | 25 | 30 | 43 | 34 |
| Total peptides | 39114 | 38677 | 38413 | 38220 | 35679 | 37870 | 37720 | 37896 |

**Table S3**. iNGNs growth factors treatment timeline after thawing a cryo-stock or splitting a maintenance plate

| Thawing/splitting day | 1 day | 2 day | 3 day | Change medium every 2 days until 80-90% confluency reached |
|---|---|---|---|---|
| E7 + Tz + TGF + FGF | E7 + TGF + FGF | skip | E7 + TGF + FGF | |

**Table S4**. iNGNs differentiation timeline treatment

| | | 2D | | 4D | | Change medium every 2 days |
|---|---|---|---|---|---|---|
| Splitting day | 1 day | 2 day | 3 day | 4 day | 5 day | |
| E7 + Tz + Dox | E7 + Dox | skip | E7 + Dox | E7/Neurobasal A = 1:1 + 2% NeuroBrew-21 | Neurobasal A + 2% NeuroBrew-21 | |

#### Organic synthesis

##### Synthesis of (*tert*-butoxycarbonyl)-*L*-tyrosine (2)[1]

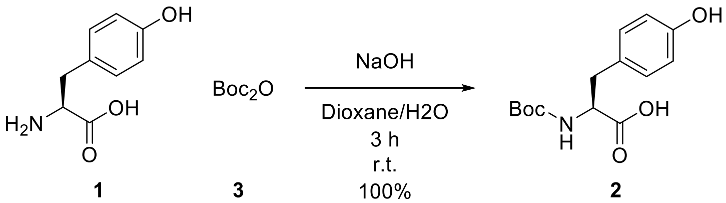

**1** **3** **2**

In a round-bottom flask, *L*-tyrosine (**1**) (11.04 mmol, 2.00 g) was dissolved in a mixture of dioxane/$H_2O$ (100 mL) in a 2/1 ratio. Afterward, alongside the addition of 1M NaOH solution (25 mL), di-*tert*-butyl dicarbonate (**3**) (12.14 mmol, 2.65 g) was added to the RM. The solution was stirred for 3 h at r.t. The RM was pre-evaporated *in vacuo,* and pH was adjusted to 2 with 6M HCl. The acidified aqueous solution was extracted with EtOAc (3 × 50 mL). Collected organic fractions were washed with brine (1 × 50 mL) and dried under anhydrous $Mg_2SO_4$. The organic solvent was evaporated *in vacuo,* and the residue was placed under high vacuum. Boc-tyrosine **2** was obtained as yellowish oily residue (3.10 g, 100%). The NMR spectra were in agreement with the literature.[2]

$^1$H NMR (400 MHz, DMSO-$d_6$) δ 12.55 (s, 1H), 9.20 (s, 1H), 7.02 (d, *J* = 8.2 Hz, 2H), 6.65 (d, *J* = 8.5 Hz, 2H), 4.02 – 3.95 (m, 1H), 2.90 – 2.83 (m, 1H), 2.73 – 2.64 (m, 1H), 1.32 (s, 9H).

##### Synthesis of prop-2-yn-1-yl (*S*)-2-amino-3-(4-(prop-2-yn-1-yloxy)phenyl)propanoate(4)[1]

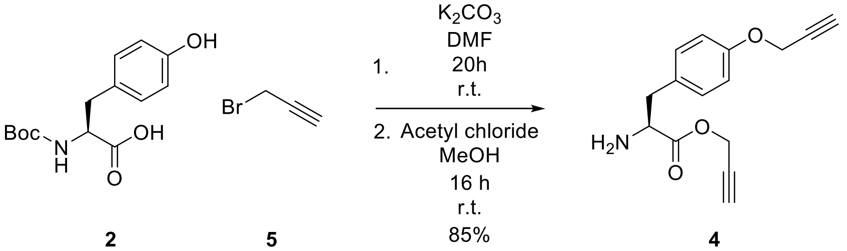

**2** **5** **4**

Under inert conditions, Boc-tyrosine **2** (5.33 mmol, 1.50 g) and $K_2CO_3$ (16.00 mmol, 2.21 g) were suspended in anhydrous DMF (40 mL). The RM was cooled in an ice bath before the dropwise addition of propargyl bromide (**5**). The resulted mixture was removed from the bath and stirred for 20 h at r.t. The RM was diluted with $H_2O$ (50 mL), and the aqueous phase was extracted with EtOAc (3 × 40 mL). The combined organic phase was washed with $H_2O$ (1 × 20 mL), brine (1 × 20 mL), and dried over anhydrous $Mg_2SO_4$. The solvent was removed *in vacuo,* and the crude residue was used directly in the next step.

To the ice-cooled anhydrous MeOH (25 mL), acetyl chloride (2.20 mL) was slowly added. The obtained solution was then added to the crude product from the previous step, and the RM was stirred overnight at r.t. The organic phase was removed *in vacuo*, and the desired product was additionally dried on a vacuum line. Propargyl ester **4** (1.17 g, 85%) was obtained as a brown solid. The NMR spectra were in agreement with the literature.[3]

$^1$H NMR (400 MHz, DMSO-$d_6$) δ 7.21 – 7.15 (m, 2H), 6.96 – 6.92 (m, 2H), 4.83 (d, *J* = 2.6 Hz, 2H), 4.78 (d, *J* = 2.4 Hz, 2H), 4.36 – 4.21 (m, 1H), 3.71 (t, *J* = 2.5 Hz, 1H), 3.59 (t, *J* = 2.4 Hz, 1H), 3.13 – 3.03 (m, 1H).

**Synthesis of (*S*)-2-amino-3-(4-(prop-2-yn-1-yloxy)phenyl)propanoic acid (Tyr-*O*-Pro, 6)**[3]

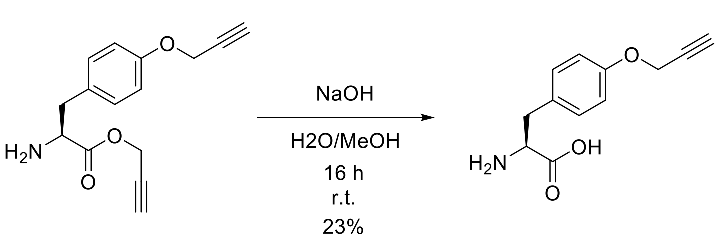

**4** **6**

Propargyl ester **4** (4.56 mmol, 1.17 g) was dissolved in MeOH (10 mL), and 1M NaOH solution (20 mL) was added to the reaction flask. After the RM was stirred for 24 h at r.t., it was acidified with conc. HCl to pH 3 and left in the fridge overnight at 4°C. The precipitated product was filtered and dried on a vacuum line. Propargyl-tyrosine **6** (0.23 g, 23%) was obtained as a brown solid. The NMR spectra were in agreement with the literature.[3]

$^1$H NMR (400 MHz, DMSO-$d_6$) δ 7.19 (d, *J* = 8.6 Hz, 2H), 6.89 (d, *J* = 8.6 Hz, 2H), 4.75 (d, *J* = 2.4 Hz, 2H), 3.56 (t, *J* = 2.3 Hz, 1H), 3.32 (dd, *J* = 8.3, 4.3 Hz, 1H), 3.07 (dd, *J* = 14.4, 4.3 Hz, 1H), 2.78 (dd, *J* = 14.4, 8.3 Hz, 1H).

**Synthesis of ONBY**

**Synthesis of *tert*-butyl *L*-tyrosinate (7)**[4]

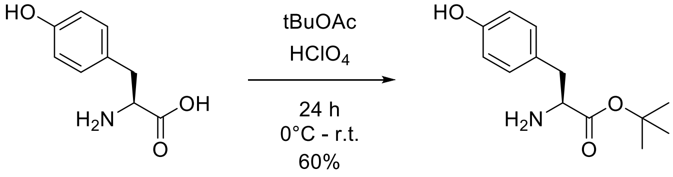

**1** **7**

Under inert conditions, perchloric acid (70%, 0.29 mL, 8.28 mmol, 1.5 eq.) was added dropwise to a suspension of *L*-tyrosine **1** (1 g, 5.52 mmol, 1 eq.) in tBuOAc (14 mL) at 0°C, and the mixture was stirred overnight at r.t. The reaction mixture was then washed with $H_2O$ (1 × 28 mL) and HCl (1 M, 1 × 14 mL). The combined aqueous phase was adjusted to pH 9 with concentrated $K_2CO_3$ solution and extracted with DCM (3 × 15 mL). The organic phase was dried over $MgSO_4$, and the solvent was removed *in vacuo*. Ester **7** (0.79 g. 60%) was obtained as a white solid. Spectra were in agreement with the literature.[5]

$^1$H NMR (400 MHz, Chloroform-*d*) δ 7.04 (d, *J* = 8.5 Hz, 2H), 6.68 (d, *J* = 8.5 Hz, 2H), 3.58 (dd, *J* = 7.8, 5.4 Hz, 1H), 2.99 (dd, *J* = 13.7, 5.4 Hz, 1H), 2.76 (dd, *J* = 13.7, 7.7 Hz, 1H), 1.45 (s, 9H).

##### Synthesis of *tert*-butyl (*tert*-butoxycarbonyl)-*L*-tyrosinate (8)

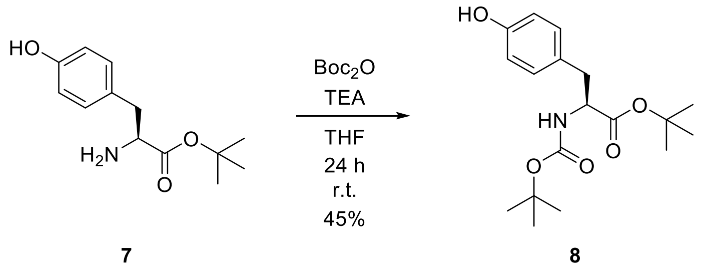

Tyr-*O*-tBu **7** (780 mg, 3.29 mmol, 1 eq.), Boc-anhydride **3** (924 mg, 3.29 mmol, 1 eq.), and triethylamine (45 µL, 0.33 mmol, 0.1 eq.) were dissolved in THF (30 mL), and the reaction mixture was stirred for 24 h at r.t. THF was then removed *in vacuo*. The mixture was dissolved in ethyl acetate (15 mL), washed with water (3 × 10 mL), and dried over $MgSO_4$. Ethyl acetate was removed *in vacuo,* and the product was purified using ethyl acetate/DCM (9:1) mixture as eluent. Boc-protected ester **8** (497 mg, 45%) was obtained as a white solid. Spectra were in agreement with the literature.[6]

$^1$H NMR (400 MHz, DMSO-*d*$_6$) δ 9.20 (s, 1H), 7.00 (d, *J* = 8.5 Hz, 2H), 6.65 (d, *J* = 8.5 Hz, 2H), 3.95 – 3.88 (m, 1H), 2.83 – 2.66 (m, 2H), 1.34 (s, 18H).

##### Synthesis of *tert*-butyl (*S*)-2-((*tert*-butoxycarbonyl)amino)-3-(4-((2-nitrobenzyl)oxy)phenyl)propanoate (9)

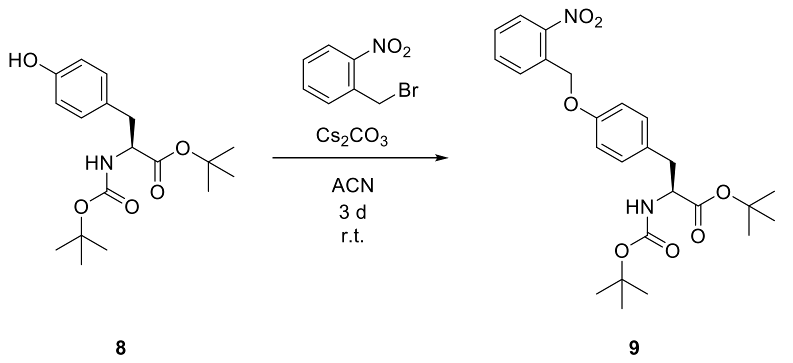

In a round-bottomed flask, Boc-Tyr-OtBu **8** (0.2 g, 0,59 mmol), 2-nitrobenzyl bromide (0.13 g, 0.59 mmol), and $Cs_2CO_3$ (0.1 g, 0.30 mmol) were dissolved in ACN (5 mL). The reaction was stirred for 3 days at r.t. The reaction mixture was filtered, and the solvent was reduced *in vacuo*. The crude product **9** was used in the subsequent step without purification and isolation.

**Synthesis of (*S*)-2-amino-3-(4-((2-nitrobenzyl)oxy)phenyl)propanoic acid (10)**

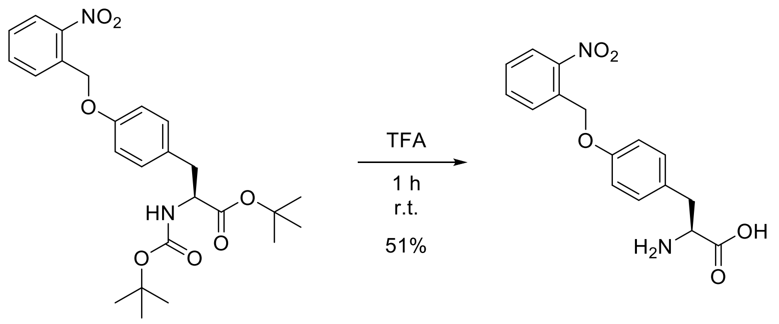

**9** **10**

The crude product from the previous step (0.28 g) was dissolved in TFA (5 mL), and the reaction mixture was mixed for 1 h at r.t. TFA was then co-evaporated with toluene *in vacuo*. Afterward, the residue was dissolved in MeOH (5 mL), and by-products were filtered off. Methanol was removed *in vacuo,* and the product was further purified by recrystallization. The solids were dissolved in 1 M NaOH, and pH was adjusted to 5 by adding 3 M HCl solution to recrystallize the product. The solution was left in the fridge overnight, and the crystals were then filtered and dried under high vacuum. The final product (96 mg, 51% yield after two reactions) was obtained as yellowish crystals. Spectra were in agreement with the literature.[7]

$^1$H NMR (400 MHz, Methanol-$d_4$) δ 8.12 (dd, *J* = 8.2, 1.3 Hz, 1H), 7.84 (dd, *J* = 7.8, 1.2 Hz, 1H), 7.72 (td, *J* = 7.6, 1.3 Hz, 1H), 7.59 – 7.52 (m, 1H), 7.20 (d, *J* = 8.6 Hz, 2H), 6.91 (d, *J* = 8.6 Hz, 2H), 5.43 (s, 2H), 3.41 (dd, *J* = 8.0, 4.8 Hz, 1H), 3.04 (dd, *J* = 13.6, 4.7 Hz, 1H), 2.73 (dd, *J* = 13.6, 8.0 Hz, 1H).

$^{13}$C NMR (101 MHz, MeOD) δ 181.78, 158.40, 148.99, 134.84, 134.75, 132.95, 131.62, 130.15, 129.76, 125.85, 115.87, 68.01, 59.01, 41.92.

MS (ESI+): *m/z* (%): 317.23 (100) $[M+Na]^+$.

### Biochemistry

#### Cell lines

##### SH-SY5Y cell line

The human neuroblastoma cell line SH-SY5Y (CRL-226) was cultivated in a high-glucose Dulbecco's Modified Eagle Medium (DMEM) that was further supplemented with 10% (v/v) heat-inactivated fetal bovine serum (FBS) and 2% (v/v) L-glutamine. Cells were maintained in cell culture dishes for adherent cells at 37°C under constant humidity and 5% $CO_2$ concentration.

##### iNGNs cell line

Before culturing the cell line, a Petri dish (p100) was coated with a Geltrex LDEV-free coating. Geltrex was added in a cold coating medium (1 eq. DMEM, 1 eq. F-12, 1% pen/strep) (10 mL) in 1/1000 dilution, mixed thoroughly, and poured directly into the dish. It was left in the incubator for at least 1 h before seeding the cells.

*E7 medium preparation*

For iNGN cells, an E7 medium was prepared. DMEM and F-12 were mixed in a 1:1 ratio, then the mixture was supplemented with L-Ala- L-Gln (2 mM), L-ascorbic acid 2-phosphate (64 mg/L), $Na_2SO_3$ (77.6 nM) and NaCl (11.2 mM). hHolo-transferrin (10 µg/mL) and hrInsulin (20 µg/mL). The E7 medium was filtered through a 0.2 µm bottle filter and stored at 4°C.

*Thawing and seeding of iNGN cells*

Freezed cells were taken out from a nitrogen tank and placed on dry ice. Cells were thawed fast by placing into 37°C bath and transferred into 50 mL falcon. Depend on the volume of the cryostock, 10 times excess of E7 medium without prewarming was added to the falcon slowly with mild agitation. Cells were subsequently pelletized by centrifuging at 200 rcf for 5 min, and the supernatant was removed. Afterward, cells were resuspended in 10 mL of prewarmed E7 media. Coating medium was removed from prewarmed dish before transferring the cell suspension dropwise. Additional growth factors and ROCK inhibitor were added in specific concentrations: TGF-β (2 ng/mL), FGF-2-IS (20 ng/mL) and Thiazovivin (**Tz**) (2 µM). The next day, the medium was changed for the fresh E7 medium adding TGF-β (2 ng/mL) and FGF-2-IS (20 ng/mL), but without addition of Tz. The medium of the same composition as for the second day was changed every two days. See **Table S3**.

*Splitting of undifferentiated iNGNs*

Cells in a maintenance plate (p100) were washed once with PBS (5 mL) before adding TrypLE™ (1 mL) and placing the plate in the incubator for 7-10 min. Tz (2 µM) was added to the prewarmed E7 medium. Supplemented with Tz, E7 medium (5 mL) was added to the plate and pipetted against its walls to form a single cell suspension. The suspension was then transferred to a 15 mL falcon and centrifuged at r.t. for 5 min at 200 rcf. The supernatant was sucked away, and the cells were suspended in a fresh prewarmed Tz-supplemented E7 medium. The volume of the medium was calculated in such a way as to reach the seeding density of 2 mil. cells for a p100 Petri dish. Before transferring the appropriate amount of cell suspension to the dish, the coating medium was discarded. Finally, cells were additionally supplemented with growth factors – TGF-β (2 ng/mL) and FGF-2-IS (20 ng/mL). The next day, the medium was exchanged for an E7 medium with growth factors but without adding Tz. The medium with growth factors was exchanged every two days. The treatment scheme is represented in **Table S3**.

*Seeding and culturing of differentiated iNGN cells*

The seeding procedure was the same as for undifferentiated cells with minor differences. The seeding density for the differentiation was 3 mil. cells per p100 dish. After preparing the cell suspension in E7 medium containing Tz (2 μM) as was previously describe, Doxycyclin (**Dox**) (0.5 μg/mL) was added instead of growth factors to initiate the differentiation. The next day, the medium was exchanged for fresh E7 without Tz but with the addition of Dox at the same concentration (0.5 μg/mL). On day four after the splitting, the medium was exchanged for a mixture of E7 and Neurobasal A in a 1:1 ratio supplemented with 2% NeuroBrew-21. After that point, the medium was exchanged for the fresh every two days. **Table S4** represents the treatment scheme.

**Cells treatment and harvesting**

SH-SY5Y or iNGN cells were treated with the stock solution of **Tyr-*O*-Pro** ($H_2O$:1M NaOH 2:1, 144 mM, sterile filtered) so that the final concentration in a dish was 0.3 mM. The incubation time for the probe was one day. Using the same volume as for the probe-treated cells, we treated the control group with a plain solvent, the same as that used to prepare the probe stock solution.

For the cells harvesting, the medium was removed, and the cells were washed once with 5 mL of a phosphate-buffered saline solution (PBS). After removing PBS from a dish, cells were scraped with 1 mL of PBS, transferred to a 1.5 mL tube, and kept shortly on ice until further manipulations. To obtain the cell pellet, the cell suspension was centrifuged at 4°C at 100 rcf. Subsequently, the supernatant was carefully removed, and the cell pellet was stored at -80°C until further use.

**Lysates preparation**

To prepare a cell lysate, the cell pellet was reconstituted in 300 μL of a lysis buffer (1% NP40, 0.2% SDS in 25 mM Hepes, 7.5 pH) by sonication with an ultrasonic tip in 1 s on/ 1 s off cycles at 20% intensity for 10 s of total time. The solution was clarified by centrifugation at 4°C at 14000 rcf for 15 min. The clear supernatant was then transferred to a new 1.5 mL tube and stored at -80°C until use

**Protein concentration measurement**

Protein concentration measurement was performed with a Pierce™ BCA Protein Assay Kit (Thermo Scientific. Cat.N. 23227).

**SP2E workflow large scale**

The enrichment samples were prepared from the lysates so that each sample contained 400 μg of proteins. Lysates were diluted to 200 μL with lysis buffer (1% NP40, 0.2% SDS in 25 mM Hepes, 7.5 pH). For each sample, 2 μL of Biotin-$N_3$ (10 mM in DMSO), 2 μL of TCEP (100 mM in $H_2O$), and 0.24 μL TBTA (83.5 mM in DMSO) were added, vortexed, spun down, and supplemented with 4 μL of $CuSO_4$ (50 mM in $H_2O$) to initiate the reaction. The reaction mixture was incubated at r.t. while shaking at 450 rpm for 1.5 h.

Streptavidin-coated magnetic beads (50 μL) were transferred to a new 1.5 mL tube and washed three times with 500 μL 0.2% SDS in PBS, sequentially vortexing and spinning down. A 1:1

mixture of hydrophobic and hydrophilic carboxylate-coated magnetic beads (100 µL) was transferred to a new 1.5 mL tube and washed three times with 500 µL MS-grade $H_2O$, vortexed, and spun down.

After completion of the click reaction, the mixture was diluted with 200 µL of 8 M Urea to a total volume of 400 µL. The reaction mixture was placed on carboxylate-coated beads, diluted with 600 µL of absolute ethanol, vortexed, and spun down. The suspension was incubated at r.t. while shaking at 950 rpm for 5 min. Afterward, the supernatant was discarded, and the beads were washed thrice with 500 µL of 80% ethanol in $H_2O$, vortexed, and spun down. After the last washing step, proteins were eluted from carboxylate-coated beads to streptavidin-coated beads. To elute the proteins, 300 µL 0.2% SDS in PBS was added to the carboxylate-coated beads, vortexed, and incubated at 40°C while shaking at 950 rpm for 5 min, and the solution was transferred to dry streptavidin-coated beads. The elution step was repeated two more times.

The eluates were incubated on streptavidin-coated beads at r.t. while shaking at 950 rpm for 20 min. The supernatant was discarded, and the beads were washed three times with 500 µL 1% NP-40 in PBS, twice with 500 µL 6 M Urea in $H_2O$, and twice with 500 µL MS-grade $H_2O$. After each round of washing, beads were vortexed and spun down.

At that point, proteins can be further processed for MS measurement or *in-gel* analysis.

**In-gel analysis of the enriched proteins**

After the last washing step, 20 µL of MS-grade $H_2O$ and 5 µL of 5 × SDS reducing Loading Buffer were added to the beads, vortexed, and spun down. The resulting suspension was incubated at 95°C while shaking at 950 rpm for 5 min. The supernatant was subsequently placed on an SDS-PAGE gel.

**Sample preparation for MS analysis**

After the last washing step, the beads were reconstituted in 80 µL of ammonium bicarbonate buffer (125 mM in $H_2O$) (ABC buffer). For the suspension, 10 µL of TCEP (100 mM) and 10 µL of chloracetamide (400 mM) solutions were added, and samples were incubated at 95°C for 5 min. After cooling the samples, trypsin (0.5 µg/µL) was added and incubated at 37°C overnight with agitation. The supernatant was transferred into a new 1.5 mL tube, and the beads were then washed with 100 µL of ABC buffer (100 mM in $H_2O$) three times. All washings were combined with the supernatant, and the resulting mixture was supplemented with 2.5 µL of formic acid. To desalt the samples, Sep-Pak C18 cartridges were used. The cartridge was flushed with 1 mL of ACN and 1 mL of ACN and FA mixture (80% + 0.5% in $H_2O$). Equilibration was performed three times with 1 mL FA (0.5% in $H_2O$). Afterward, the sample was loaded into the cartridge. The column was then washed three times with 1 mL of FA (0.5% in $H_2O$). Peptides were eluted with ACN and FA mixture (80% + 0.5% in $H_2O$) in two 250 µL batches. The samples were dried in a SpeedVac.

Dry peptides were then reconstituted in 30 µL FA (1% in $H_2O$), vortexed, and placed in a sonication bath for 15 min. Samples were afterward spun down and transferred into MS vials.

**SP2E workflow small scale**

Lysates containing 100 µg of proteins were diluted to 19 µL with lysis buffer (1% NP40, 0.2% SDS in 25 mM Hepes, 7.5 pH). For each sample, 0.2 µL Biotin-$N_3$ (10 mM in DMSO), 0.2 µL of TCEP

(100 mM in $H_2O$), and 0.125 µL TBTA (16.7 mM in DMSO) was added, vortexed, spun down, and supplemented with 0.4 µL of $CuSO_4$ (50 mM in $H_2O$) to initiate the reaction. The reaction mixture was incubated at r.t. while shaking at 450 rpm for 1.5 h.

After completion of the click reaction, each sample was diluted with 60 µL of 8M urea. A 1:1 mixture of hydrophobic and hydrophilic carboxylate-coated magnetic beads (100 µL) was washed three times with 100 µL MS-grade $H_2O$, and the reaction mixture was placed on the beads, diluted with 100 µL of absolute ethanol and vortexed. The suspension was incubated at r.t. while shaking at 950 rpm for 5 min. Afterward, the supernatant was discarded, and the beads were washed thrice with 150 µL of 80% ethanol in $H_2O$ and once with 150 µL acetonitrile (LC-MS). Proteins were eluted separately by adding 60 µL of 0.2 % SDS in PBS. For this, beads were resuspended and incubated for 5 min at 40 ° C and 950 rpm. The supernatant was directly transferred onto 50 µL equilibrated streptavidin-coated magnetic beads (3 times prewashed with 100 µL 0.2% SDS in PBS). The elution step was repeated twice and the combined beads mixture was incubated at r.t. while shaking at 800 rpm for 1 h.

The supernatant was discarded, and the beads were washed three times with 150 µL 1% NP-40 in PBS, twice with 150 µL 6 M Urea in $H_2O$, and twice with 500 µL MS-grade $H_2O$. After each round of washing, the beads were incubated at r.t. while shaking at 800 rpm for 1 min. The rinsed beads mixtures were resuspended in 50 µL 50 mM TEAB, and the proteins were digested overnight at 37°C by adding 1.5 µL sequencing grade trypsin (0.5 mg/mL). The following day, the beads were washed twice with 20 µL of 50 mM TEAB buffer and twice with 20 µL 0.5% FA, and the wash fractions were collected and combined. The beads were incubated for 5 min at 40°C and 600 rpm for each washing step. The combined washed fractions were acidified by adding 0.9 µL formic acid (FA) and transferred to MS vials.

**In-gel analysis**

The resolution of proteins was made with the SDS-PAGE method using 10% acrylamide gels. Before loading onto the gel, a protein solution (20 µL) was mixed with 5 × SDS reducing loading buffer (5 µL) (10% (w/v) SDS, 50% (v/v) glycerol, 25% (v/v) β-mercaptoethanol, 0.5% (w/v) bromphenol blue, 315 mM Tris/HCl, pH 6.8) and placed in wells. As a reference, two types of protein markers were used: BenchMark™ Fluorescent Protein Standard (Invitrogen™), Color Prestained Protein Standard, Broad Range (10-250 kDa) (New England Biolabs GmbH). Afterward, the gel was scanned on Amersham Imager 680 (GE Healthcare).

**Western blot**

After separating the proteins on a gel, they were blotted on a PVDF membrane using a Semi-Dry Blotter (Bio-Rad). Before making a blotting sandwich, a thick blot paper was soaked in a blot buffer (48 mM Tris, 39 mM glycine, 0.0375% (m/v) SDS, 20% (v/v) methanol) for 5 min, and a membrane was incubated in methanol. After the blotting, the membrane was set in a blocking solution (0.5 g nonfat dried milk powder in 10 mL PBST (PBS + 0.5% Tween)) for 60 min to hide all nonspecific binding sites. The membrane was then placed in the primary antibody of interest solution and incubated at 4°C overnight. The next day, the membrane was washed 3 × 10 min with PBST solution before incubation with the secondary HRP-linked antibody solution at r.t. for 1 h. The membrane was then washed with PBST 3 × 10 min. Before scanning the membrane on Amersham Imager 680 (GE Healthcare), it was wetted with the ECL substrate and the peroxide solution in a 1:1 ratio.

**Whole proteome samples preparation**

SH-SY cell lysates were prepared under standard conditions using standard treatment and harvesting procedures. The volume of a lysate containing 100 µg of proteins was normalized to 400 µL with lysis buffer. A mixture of hydrophilic and hydrophobic carboxylate-coated magnetic beads was washed thrice with 500 µL of MS-grade $H_2O$. The sample was added to the beads and thoroughly mixed. 600 µL of absolute EtOH was added, and the mixture was incubated at r.t. for 5 min with agitation. Subsequently, the beads were washed with EtOH (80% in $H_2O$) three times.

To cleave the proteins, on-beads digestion was performed. The procedure was identical to those which was used for MS-samples preparation. In brief, beads were reconstituted in ABC buffer alongside reducing and alkylating agent and boiled. After the samples were cooled down, trypsin was added and left overnight at 37°C. The supernatant was then placed in a new 1.5 mL tube, and the beads were washed several times with ABC buffer. After desalting the peptide mixture on a C18 column, samples were dried on a SpeedVac.

The dry peptides were reconstituted in 200 µL of FA (1% in $H_2O$) and transferred into MS vials.

**Enrichments with trifunctional linker (5/6-TAMRA-$N_3$-biotin)**

Lysates from iNGNs were prepared on the 2$^{nd}$ (**2D**) and 4$^{th}$ (**4D**) day after starting of the differentiation using standard protocols. Cells were treated with 0.3 mM of the probe and incubating for 1 day before harvesting. Protein concentration in the lysates were evaluated with Pierce™ BCA Protein Assay Kit. Large-scale SP2E protocol was implemented as a template for the preparation of the probes with minor differences. Instead of biotin-$N_3$ probe, 5/6-TAMRA-$N_3$-biotin (trifunctional linker) probe underwent the click reaction with the same parameters as in the original SP2E protocol. After enrichment, proteins were eluted from the beads into separate 1.5 mL tube instead of been digested with trypsin. For the elution, 20 µL of $H_2O$, and 5 µL of 5 × SDS reducing Loading Buffer were placed on beads, vortexed and incubated at 95°C at 850 rpm for 5 min. Afterward, samples were placed on a magnet rack and let cooled at r.t. Supernatants were further used for *in-gel* analysis. For WB analysis, rat anti-mapre1 (Abcam - ab53358) antibody was used at 2 µg/mL concentration. A goat anti-rat HRP-linked secondary antibody (Cell Signaling Technology - 7077S) was used in 1/1000 dilution.

**Tubulin fraction isolation**

SH-SY5Y cells were grown in a p150 Petri dish in DMEM under standard conditions and with the standard probe treatment scheme. Cell pellets were prepared with a standard protocol and placed in 1.3 mL Hepes/Pipes buffer (25 mM Hepes, 60 mM Pipes) without the addition of EGTA. The cell suspension was sonicated on ice with a microsonic tip at 50% intensity for 1 min in 10 s on/off cycles. The samples were then centrifuged at 20k rcf at 4°C for 1 h. The first pellet was kept to confirm the absence of tubulins. The supernatant was transferred to a new 1.5 mL tube and additional additives were added – $MgCl_2$ (3 mM final concentration), Taxol (50 µM final concentration), and GTP (2 mM final concentration). The samples were then incubated at 37°C for 1 h. After incubation, the supernatants were placed on 0.2 mL sucrose cushion containing 20 µM Taxol and 1.5 mM GTP, and the samples were ultracentrifuged at 150k rfc at 37°C for 2 h. After the centrifugation, MTs pellet was then resuspended in 0.25 mL of a click buffer (1% NP40, 0.2% SDS in 25 mM Hepes, 7.5 pH) and sonicated on ice with a microsonic tip at 20% intensity for 10 s in 1 s on/off cycles. The supernatant was additionally clarified by centrifugation at 10k rcf at 4°C for 10 m. Protein concentration in the lysates from two fraction were evaluated with Pierce™ BCA Protein Assay Kit. Afterward, click reaction was performed with a fraction containing cell debris (PI – first pellet), and MTs fraction (PII – second pellet). For the click reaction, 50 µg of proteins in 100 µL of total volume were prepared. To each sample, 1 µL TAMRA-$N_3$ (10 mM in DMSO), 1 µL of TCEP (100 mM in $H_2O$), and 0.12 µL TBTA (83.5 mM in DMSO) was added, vortexed, spun down, and supplemented with 2 µL of $CuSO_4$ (50 mM in $H_2O$) to initiate the reaction. The reaction mixture was incubated at r.t. while shaking at 450 rpm for 1.5 h. After the click reaction, proteins were precipitated in acetone ON. Next, they were centrifuged at 14k rcf at 4°C for 10 min. The pellets were then washed twice with cold methanol. After the last wash, pellets were reconstituted in 50 µL $H_2O$. To the new tube, 10 µL of the solution was transferred and mixed with 2 µL of 5 × SDS reducing Loading Buffer, and incubated at 95°C for 5 min with agitation. The resulting mixture was loaded on a 10% acrylamide gel. Subsequently, proteins were blotted and stained with a rat anti-alpha-tubulin antibody (MA1-80017, Invitrogen) at 2 µg/mL concentration.

**Fluorescence microscopy**

The SH-SY5Y neuroblastoma cell line was cultivated in slides - Nunc™ Lab-Tek™ II Chamber Slide™ System (Thermo Scientific™) to prepare samples for fluorescence microscopy. The seeding density for each well was $2 * 10^4$ cells/mL, the medium volume for one well was 0.5 mL. After one day of cell growth, the cultivating medium was exchanged for the fresh medium containing the propargyl-tyrosine probe at a concentration of 0.2 mM. The cultivation medium was exchanged for the medium containing 1% DMSO in a control group. The next day, the cells were washed trice with 0.5 mL of PBS before the fixation step. Cells were incubated in 0.5 mL of 4% PFA for 15 min with gentle mixing to fix the cells. Then, cells were washed trice with 0.5 mL PBS before permeating in 0.5 mL 0.1% Triton X-100 in PBS for 15 min with gentle agitation. After the permeabilization, cells were washed trice with PBS. Blocking nonspecific binding sites was performed in 0.5 mL of 1% BSA in PBS solution for 1 h. Subsequently, the washing step was performed.

After preparing the cells for the click reaction, the Master Mix was prepared: $Cu_2SO_4$ and TBTA solutions were mixed so that the final concentration after dilution was 1 mM and 5 µM, respectively. To the solution, sodium ascorbate was added to reach the final concentration of 10 mM. The mixture was incubated at r.t. for 15 min. TAMRA-azide was added in 10 µM final concentration. The cells were incubated with the resulting solution at r.t. for 1.5 h in the dark under mild agitation. Subsequently, cells were thoroughly washed with 0.5 mL of PBS for 15 min trice to remove all unbounded TAMRA. Cells were incubated at 4°C overnight in a blocking solution treated with primary antibodies in 1/1000 dilution. Rabbit anti-MAP2 polyclonal antibody (PA5-110744, Thermo Fisher) and rat anti-tubulin monoclonal antibody (MA1-80017, Thermo Fisher) were used. After trice washing with PBS, cells were incubated with secondary fluorophore-labeled antibodies in 1/500 dilution at r.t. for 1 h (anti-Rabbit IgG (H+L) F(ab')2 Fragment (Alexa Fluor (R) 488 conjugate) (4412S, Cell Signal) and anti-Rat IgG (H+L) Cross-Adsorbed Secondary Antibody (Alexa Fluor 647) (A-21247, Thermo Fisher)). As the last step, anti-fade fluorescence mounting medium (Fluoroshield (ab104135, abcam)) was applied to slides. Images were obtained on a Leica confocal microscope.

**Cytotoxicity measurement**

Cells were seeded in triplicates for each concentration in a transparent flat-bottomed 96-well plate at a density of 5000 cells per well in a total 100 µL and left to settle down overnight. Next, the medium was exchanged for a fresh medium supplemented with the probe to reach concentrations in a range of 0 to 2000 µM in a well, while the control samples were treated with a medium containing 1% DMSO. The cells were incubated with the probe for another 24 h. In each well, 20 µL of 3-(4,5-dimethylthiazol-2-yl)-2,5-diphenyltetrazolium bromide (**MTT**) solution was added and incubated for 4 h. Subsequently, the medium was removed, and the cells were lysed by adding 200 µL of DMSO. Absorbance was measured at 570 nm wavelength using 630 nm as a reference.

**Competition experiment**

The human neuroblastoma cell line SH-SY5Y was cultured under standard conditions in five dishes. The **Tyr-*O*-Pro** probe concentration was maintained at a constant concentration of 0.3 mM in all probe-treated plates. The competitor's concentration was changed in tenfold steps compared to the probe concentration of interest – 0.03 mM, 0.3 mM, and 3 mM. Two dishes were prepared as a control group, where one of them contained cells without any treatment (negative control), and another was treated only with the probe (0.3 mM) without the competitor (positive control). The incubation time was set to 1 day, after which the cells were harvested and processed under standard conditions as described above. Briefly, after collecting cells and measuring protein concentration, samples underwent click reaction with TAMRA-azide, proteins were precipitated in acetone overnight and separated by SDS-Page. Gel images were taken by Amersham Imager.

**CHX inhibition experiment**

The human neuroblastoma cell line SH-SY5Y was cultured in three plates under standard conditions. One plate served as a negative control without any supplement. The second plate was treated only with the **Tyr-*O*-Pro** probe (0.3 mM). The experiment plate was first treated with cycloheximide (50 µg/mL) 30 min prior addition of the **Tyr-*O*-Pro** probe (0.3 mM). After one day of incubation, cells were harvested and protein concentration was standardly measured. From each plate, 150 ug of proteins in 150 uL of buffer were then subjected to click reaction with TAMRA-azide under standard conditions as described above. After 1.5 h, 20 µL of the reaction mixture was mixed with 5 µL of 5 x SDS loading buffer, and proteins were separated by SDS-Page. The gels were imaged with an Amersham imager.

**MS measurement**

MS measurements were performed on an Orbitrap Eclipse Tribrid Mass Spectrometer (Thermo Fisher Scientific) coupled to an UltiMate 3000 Nano-HPLC (Thermo Fisher Scientific) via an EASY-Spray source (Thermo Fisher Scientific) and FAIMS interface (Thermo Fisher Scientific). First, peptides were loaded on an Acclaim PepMap 100 µ-precolumn cartridge (5 µm, 100 Å, 300 µm ID x 5 mm, Thermo Fisher Scientific). Then, peptides were separated at 40°C on a PicoTip emitter (noncoated, 15 cm, 75 µm ID, 8 µm tip, New Objective) that was *in-house* packed with Reprosil-Pur 120 C18-AQ material (1.9 µm, 150 Å, Dr. A. Maisch GmbH).

Buffer composition. Buffer A consist of MS-grade $H_2O$ supplemented with 0.1% FA. Buffer B consists of acetonitrile supplemented with 0.1% FA.

Following LC gradient was used for the short acquisition method: 4% buffer B (minutes 0 - 5), 4% - 7% buffer B (minutes 5 - 6), 7% - 24.8% (minutes 6 - 36), 24.8% - 35.2% buffer B (minutes 36 -

41), 35.2% - 80% buffer B (minutes 41 – 41,1), 80% buffer B (minutes 41.1 – 46), 80% - 4% buffer B (minutes 46 – 46.1) then hold on 4% until minute 60. The flow rate was 0.3 uL/min.

Following LC gradient was used for the long acquisition method: 4% buffer B (minutes 0 - 5), 4% - 7% buffer B (minutes 5 - 6), 7% - 24.8% buffer B (minutes 6 – 105), 24.8% - 35.2% buffer B (minutes 105 – 126), 35.2% - 80% buffer B (minutes 126 – 126.1), 80% buffer B (minutes 126.1 – 131), 80% - 4% buffer B (minutes 130 – 130.1) then hold on 4% until minute 150. The flow rate was 0.3 uL/min.

*Data independent acquisition*

The DIA duty cycle was consisted of one MS1 scan followed by 30 MS2 scans with an isolation window of the 4 m/z range, overlapping with an adjacent window at the 2 m/z range. MS1 scan was conducted with Orbitrap at 60000 resolution power and a scan range of 200 – 1800 m/z with an adjusted RF lens at 30%. MS2 scans were conducted with Orbitrap at 30000 resolution power, RF lens was set to 30%. The precursor mass window was restricted to a 500 – 740 m/z range. HCD fragmentation was enabled as an activation method with a fixed collision energy of 35%. FAIMS was performed with one alternating CV at -45V for both MS1 and MS2 scans during the duty cycle.

*Data dependent acquisition*

For measurements of DDA-MS2 mode, the Orbitrap Eclipse Tribrid Mass Spectrometer was operated with the following settings: Polarity: positive; MS1 resolution: 240k; MS1 AGC target: standard; MS1 maximum injection time: 50 ms; MS1 scan range: m/z 375-1500; MS2 ion trap scan rate: rapid; MS2 AGC target: standard; MS2 maximum injection time: 35 ms; MS2 cycle time: 1.7 s; MS2 isolation window: m/z 1.2; HCD stepped normalised collision energy: 30%; intensity threshold: 1.0e4 counts; included charge states: 2-6; dynamic exclusion: 60 s. FAIMS was performed with two alternating CVs including -50 V and -70 V.

**Quantification and statistical analysis**

*Computational evaluation of DIA raw files*

Raw files were converted in the first step with “MSConvertGUI” as a part of the “ProteoWizard” software package (http://www.proteowizard.org/download.html) to an output mzML format applying the “peakPicking” filter with “vendor msLevel=1”, and the “Demultiplex” filter with parameters “Overlap Only” and “mass error” set to 10 ppm.

*Standalone DIA-NN software under version 1.8.1 was used for protein identification and quantification*.

First, a spectral library was predicted *in silico* by the software’s deep learning-based spectra, RTs and IMs prediction using Uniprot *H. sapiens* decoyed FASTA (canonical and isoforms – May 2022). FASTA digest for library-free search/library generation option was enabled for this. Spectral library prediction was performed in 4 batches of 10 samples each to decrease computational load.

Second, all samples (40) were processed together without spectral library generation, with a match between runs (MBR) option and precursor FDR level set at 1%. Previously generated spectral libraries were implemented during the search by presenting the command (‘--lib [file name]’) into the command box.

DIA-NN search settings: Library generation was set to smart profiling, Quantification strategy - Robust LC. The mass accuracy the MS1 accuracy, and the scan window were set to 0 to allow the software to identify optimal conditions. The precursor m/z range was changed to 500-740 m/z to fit the measuring parameters.

Carbamidomethylation was set as a fixed modification, oxidation of methionine and N-term acetylation were set as variable modifications. On the contrary, the small-scale samples of the 96-well plate were calculated without carbamidomethylation as a fixed modification.

Statistical analysis of the DIA-NN result table “report.pg_matrix.csv” was done with Perseus 1.6.10.43.[8] First, potential contaminants, as well as reverse peptides, were removed from the table. Then the LFQ intensities were $log_2$-transformed. Afterward, the rows corresponding to a time point were divided into two groups – Control and Probe-treated sample. Subsequently, the groups were filtered for at least three valid values out of four rows in at least one group, and the missing values were replaced from a normal distribution with the downshift of 1.8. The $-log_{10}$(p-values) were obtained by a two-sided one-sample Student’s t-test over replicates with the initial significance level of $p = 0.05$. Fold change values, as well as p-values, were obtained for each time point.

*Computational evaluation of DDA raw files*

Raw files were converted in the first step, with “MSConvertGUI” as a part of the “ProteoWizard” software package (http://www.proteowizard.org/download.html) to an output mzML format applying the “peakPicking” filter with “vendor msLevel=1”. Converted files were further calculated with proteomics pipeline FragPipe version 18.0 containing a search engine MSFragger version 3.5. Search settings were established as follows: closed search approach with precursor mass tolerance in a range -20 – 20 ppm and fragment mass tolerance 20 ppm. Carbamidomethylation was set as a fixed modification and methionine oxidation as well as N-terminal acetylation as a variable modification. As a variable modification, a mass delta of 38.0156 m/z was set corresponding to a propargyl modification, occurring on Y. False discovery rate determination was carried out using a decoy database and thresholds were set to 1% FDR both at a peptide-spectrum match and at protein levels.

Mass spectrometry-based proteomics data have been deposited at ProteomeXchange. The accession number is PXD035044. The access is available using following credentials: and 7bN0xiKI.

### NMR Spectra

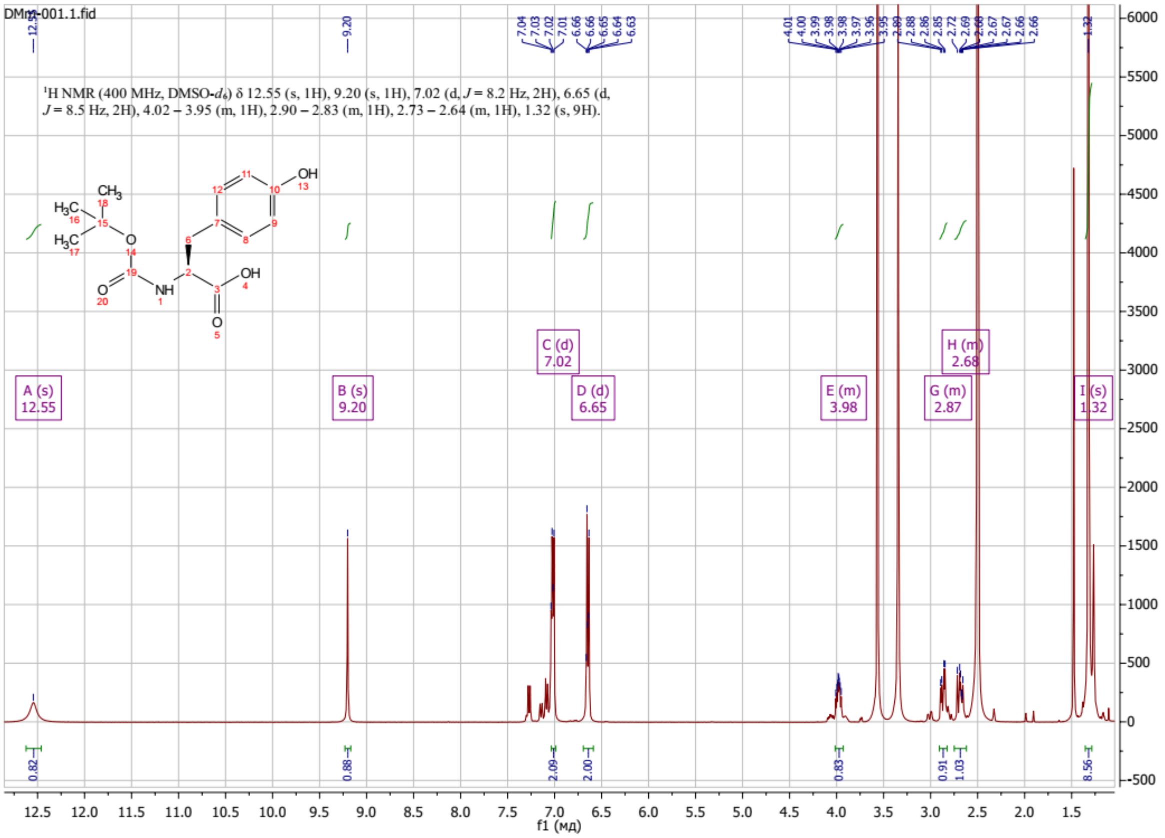
DMm-001.1.fid
$^1$H NMR (400 MHz, DMSO-$d_6$) δ 12.55 (s, 1H), 9.20 (s, 1H), 7.02 (d, $J$ = 8.2 Hz, 2H), 6.65 (d, $J$ = 8.5 Hz, 2H), 4.02 – 3.95 (m, 1H), 2.90 – 2.83 (m, 1H), 2.73 – 2.64 (m, 1H), 1.32 (s, 9H).
A (s) 12.55
B (s) 9.20
C (d) 7.02
D (d) 6.65
E (m) 3.98
G (m) 2.87
H (m) 2.68
I (s) 1.32
f1 (мд)

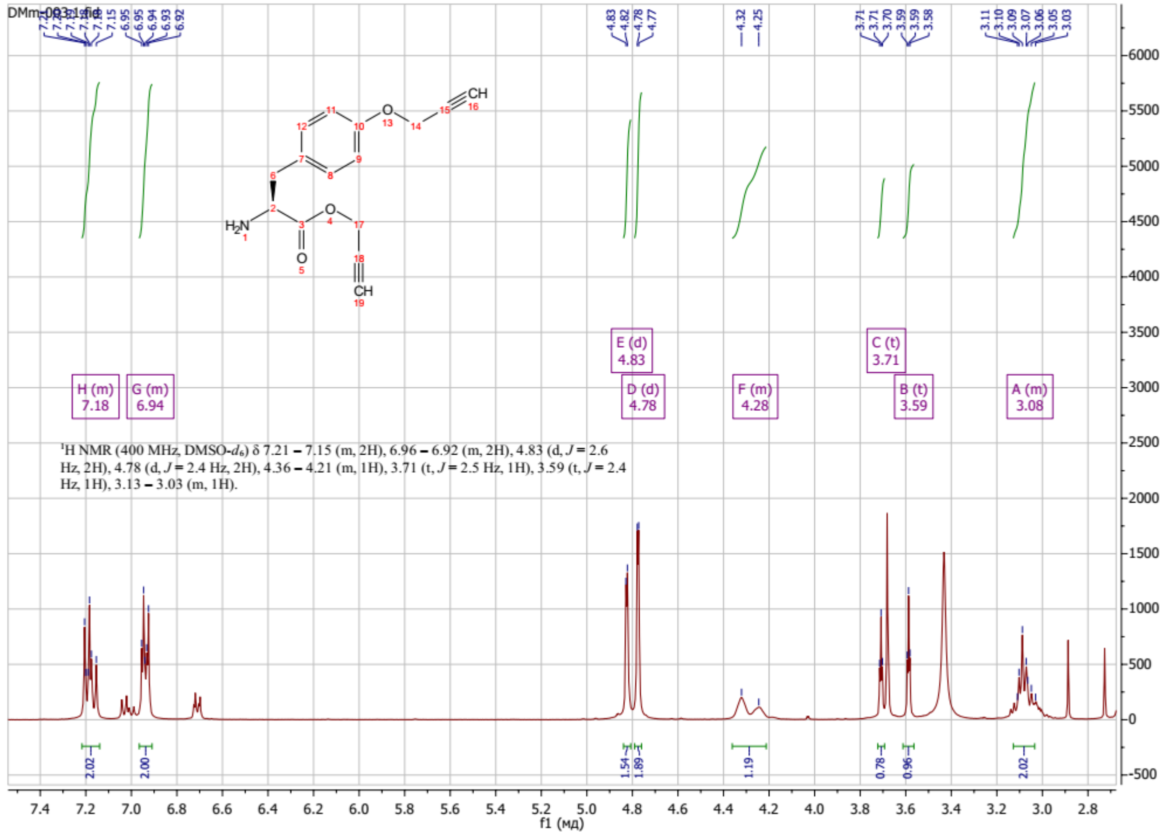
$^1$H NMR (400 MHz, DMSO-$d_6$) δ 7.21 – 7.15 (m, 2H), 6.96 – 6.92 (m, 2H), 4.83 (d, $J$ = 2.6 Hz, 2H), 4.78 (d, $J$ = 2.4 Hz, 2H), 4.36 – 4.21 (m, 1H), 3.71 (t, $J$ = 2.5 Hz, 1H), 3.59 (t, $J$ = 2.4 Hz, 1H), 3.13 – 3.03 (m, 1H).
H (m) 7.18
G (m) 6.94
E (d) 4.83
D (d) 4.78
F (m) 4.28
C (t) 3.71
B (t) 3.59
A (m) 3.08
f1 (мд)

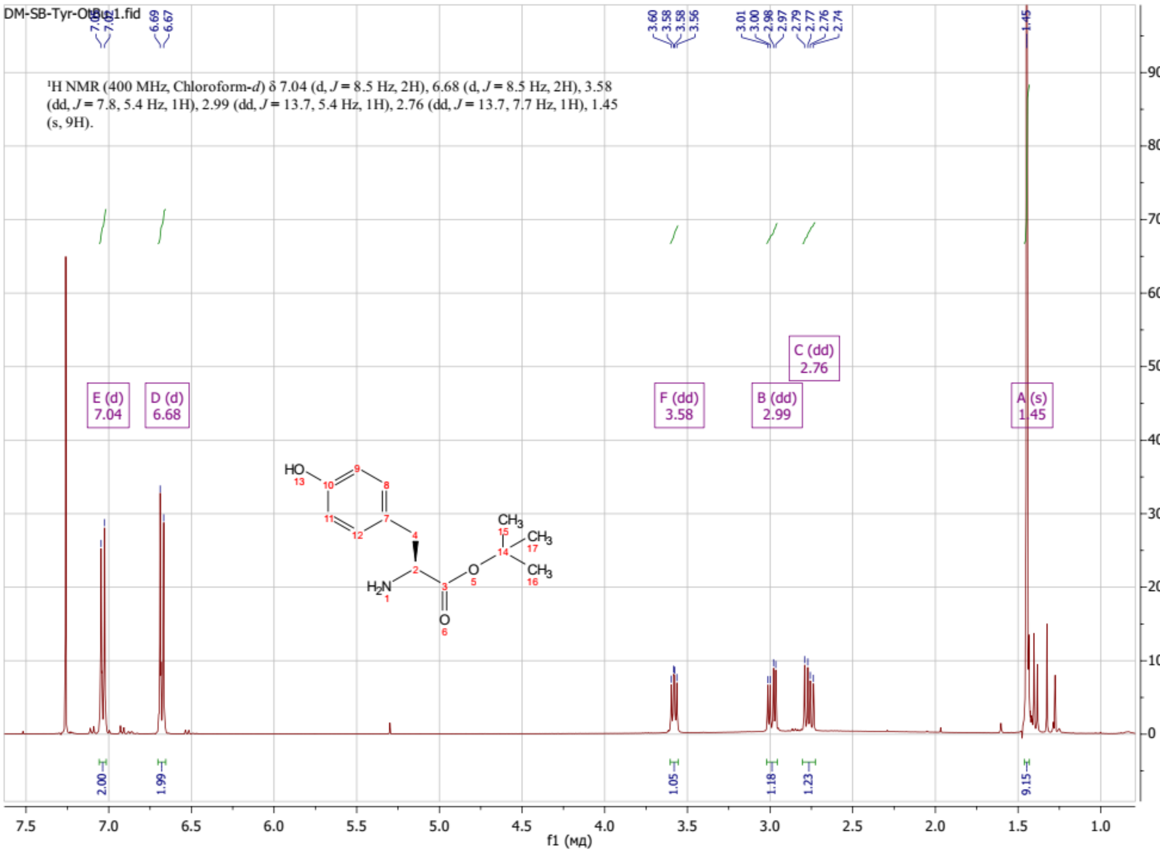
DM-SB-Tyr-OtBu.1.fid
¹H NMR (400 MHz, Chloroform-*d*) δ 7.04 (d, *J* = 8.5 Hz, 2H), 6.68 (d, *J* = 8.5 Hz, 2H), 3.58 (dd, *J* = 7.8, 5.4 Hz, 1H), 2.99 (dd, *J* = 13.7, 5.4 Hz, 1H), 2.76 (dd, *J* = 13.7, 7.7 Hz, 1H), 1.45 (s, 9H).
E (d) 7.04
D (d) 6.68
F (dd) 3.58
B (dd) 2.99
C (dd) 2.76
A (s) 1.45
f1 (мд)

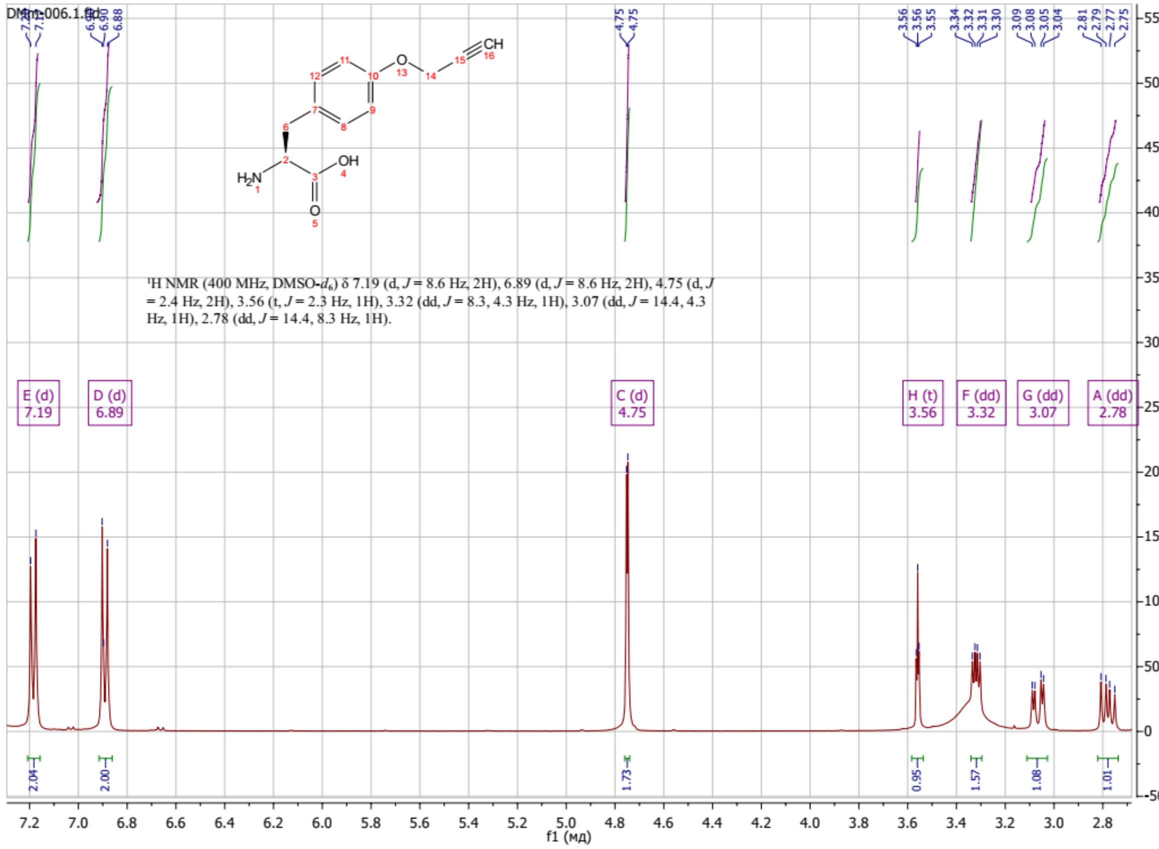
DMm-006.1.fid
¹H NMR (400 MHz, DMSO-*d*₆) δ 7.19 (d, *J* = 8.6 Hz, 2H), 6.89 (d, *J* = 8.6 Hz, 2H), 4.75 (d, *J* = 2.4 Hz, 2H), 3.56 (t, *J* = 2.3 Hz, 1H), 3.32 (dd, *J* = 8.3, 4.3 Hz, 1H), 3.07 (dd, *J* = 14.4, 4.3 Hz, 1H), 2.78 (dd, *J* = 14.4, 8.3 Hz, 1H).
E (d) 7.19
D (d) 6.89
C (d) 4.75
H (t) 3.56
F (dd) 3.32
G (dd) 3.07
A (dd) 2.78
f1 (мд)

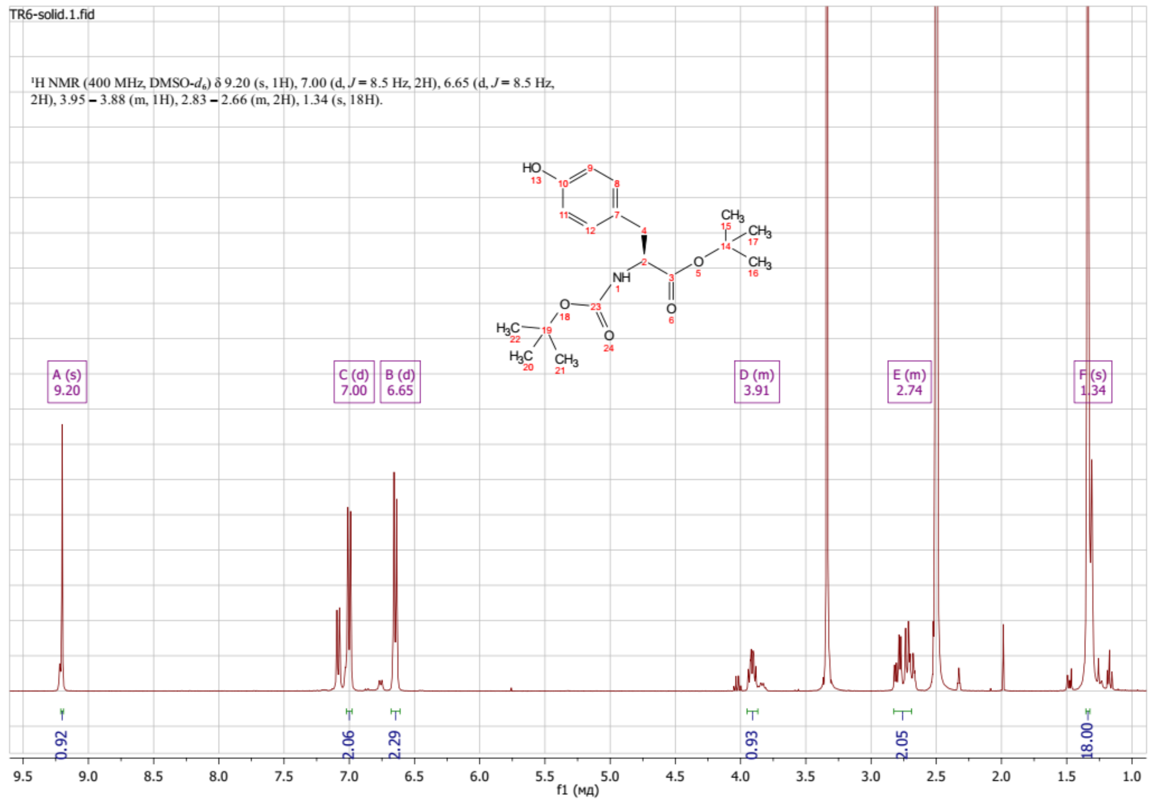
TR6-solid.1.fid
¹H NMR (400 MHz, DMSO-$d_6$) δ 9.20 (s, 1H), 7.00 (d, $J$ = 8.5 Hz, 2H), 6.65 (d, $J$ = 8.5 Hz, 2H), 3.95 – 3.88 (m, 1H), 2.83 – 2.66 (m, 2H), 1.34 (s, 18H).
A (s) 9.20
C (d) 7.00
B (d) 6.65
D (m) 3.91
E (m) 2.74
F (s) 1.34
0.92
2.06
2.29
0.93
2.05
18.00
f1 (мд)

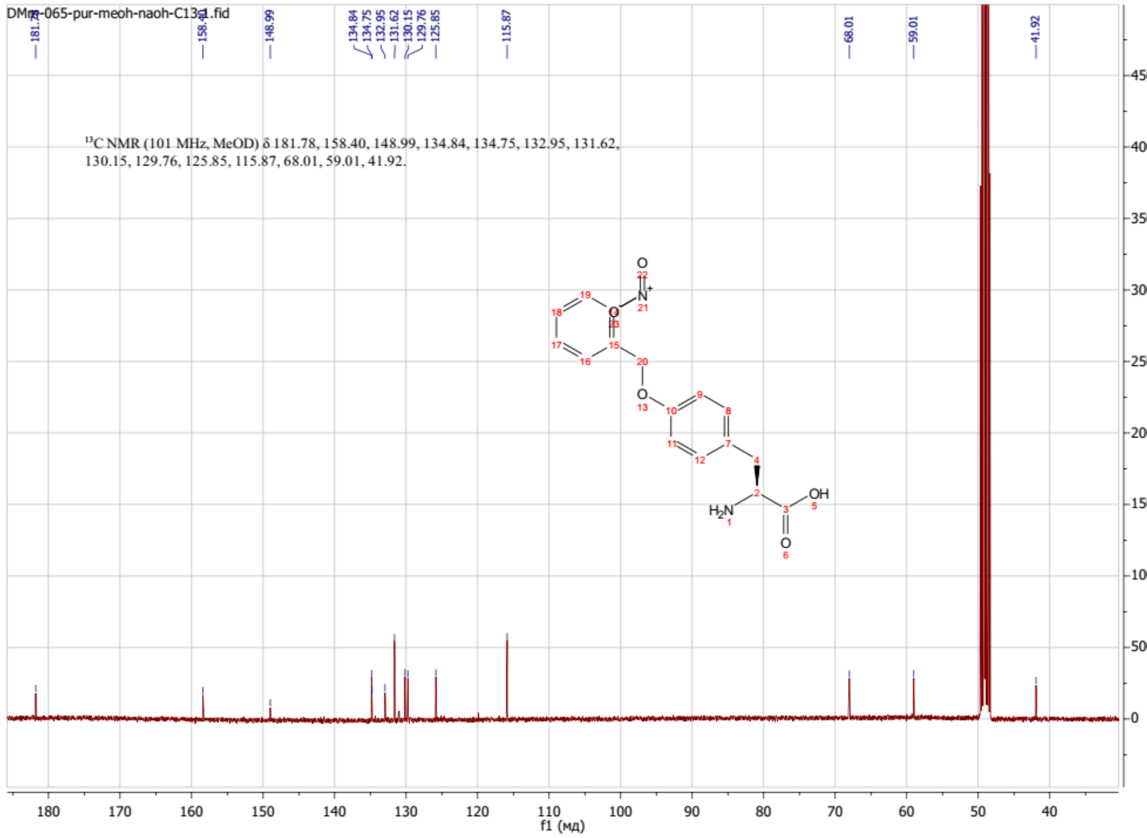
DMm-065-pur-meoh-naoh-C13-1.fid
¹³C NMR (101 MHz, MeOD) δ 181.78, 158.40, 148.99, 134.84, 134.75, 132.95, 131.62, 130.15, 129.76, 125.85, 115.87, 68.01, 59.01, 41.92.
148.99
134.84
134.75
132.95
131.62
130.15
129.76
125.85
115.87
68.01
59.01
41.92
f1 (мд)

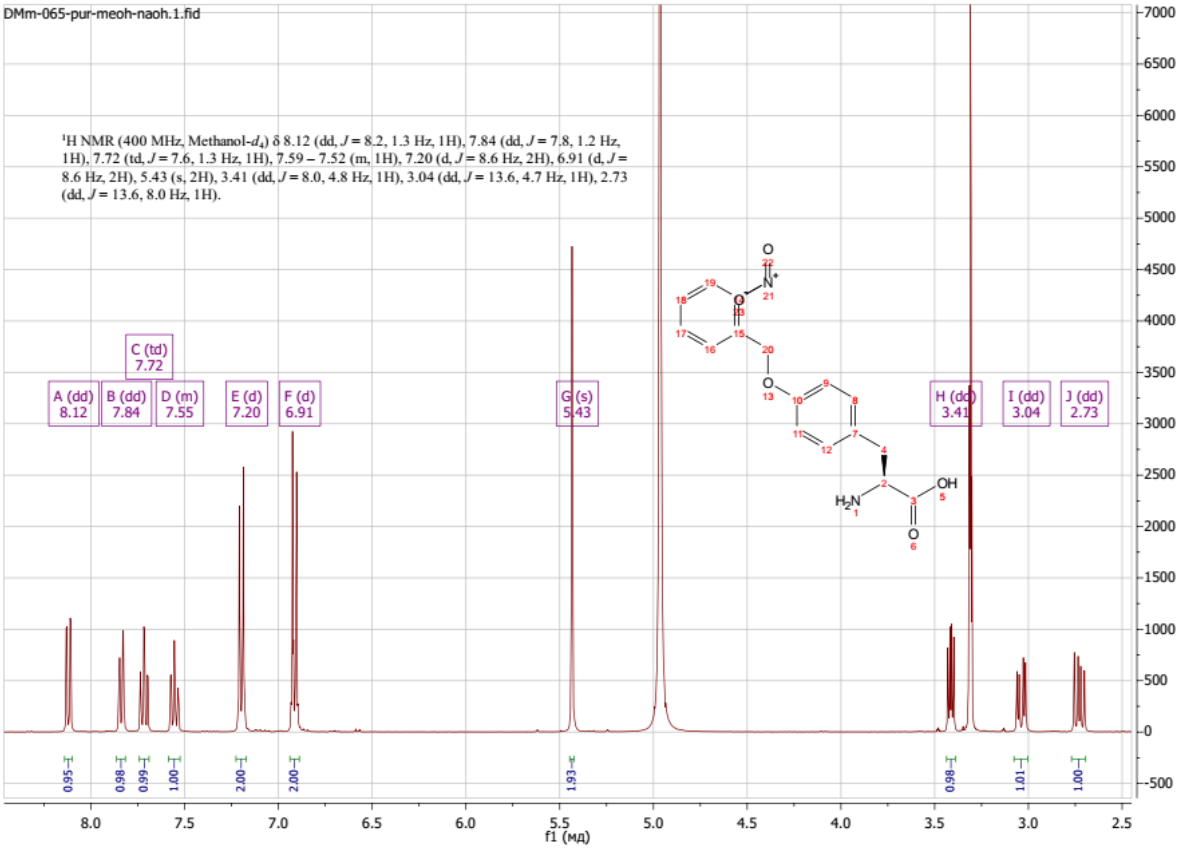
DMm-065-pur-meoh-naoh.1.fid
1H NMR (400 MHz, Methanol-$d_4$) δ 8.12 (dd, J = 8.2, 1.3 Hz, 1H), 7.84 (dd, J = 7.8, 1.2 Hz, 1H), 7.72 (td, J = 7.6, 1.3 Hz, 1H), 7.59 – 7.52 (m, 1H), 7.20 (d, J = 8.6 Hz, 2H), 6.91 (d, J = 8.6 Hz, 2H), 5.43 (s, 2H), 3.41 (dd, J = 8.0, 4.8 Hz, 1H), 3.04 (dd, J = 13.6, 4.7 Hz, 1H), 2.73 (dd, J = 13.6, 8.0 Hz, 1H).
A (dd) 8.12
B (dd) 7.84
C (td) 7.72
D (m) 7.55
E (d) 7.20
F (d) 6.91
G (s) 5.43
H (dd) 3.41
I (dd) 3.04
J (dd) 2.73
0.95
0.98
0.99
1.00
2.00
2.00
1.93
0.98
1.01
1.00
f1 (мд)
